# Screening for pro-adhesive compounds and their relevance as therapeutic approach in Arrhythmogenic Cardiomyopathy

**DOI:** 10.1101/2025.03.23.643549

**Authors:** Pauline Hanns, Robin Colpaert, Ramon Castellanos-Martinez, Frauke Weidner, Stefanie Beensen, Franziska Matthias, Lifen Xu, Gabriela M. Kuster, Camilla Schinner

## Abstract

**Background:** Arrhythmogenic Cardiomyopathy (ACM) is one of the major causes of sudden cardiac death in young adults. With the underlying patho-mechanisms not well understood, current therapeutic approaches for this genetic disease are solely symptomatic. A recent study demonstrates that loss of cell-cell adhesion is an important initial step leading to ACM. Because loss of cell-cell adhesion is considered a key initial step, we aim to identify new compounds from a drug library, which can restore intercellular adhesion and potentially serve as therapeutics for ACM.

**Methods:** We established a 2D cell adhesion-based high-throughput platform and screened an FDA-approved drug library. To model loss of intercellular adhesion, human cell lines deficient for the desmosomal adhesion molecule desmoglein-2 (DSG2) were employed and the revealed top candidates were validated in cells expressing different ACM patient mutations. The therapeutic potential of the top hit dexamethasone was evaluated in an inducible ACM disease mouse model using ECG, echocardiography and histology. Phospho-proteomic analysis was applied to investigate protective drug mechanisms.

**Results:** We established a set-up to sensitively detect changes in cell-cell adhesion via a high-throughput platform and applied this approach to identify a large pool of adhesion-strengthening compounds in DSG2 deficient cells. The pro-adhesive effect of selected candidates was validated for different ACM patient mutations inducing defective cell cohesion. Importantly, *in vivo* administration of the selected top pro-adhesive drug dexamethasone rescued impaired right ventricular function in an ACM mouse model. Phospho-proteome analysis suggests modulation of AKT1/AMPKα1 signaling and changes in cardiac contractility and junctional components as factors contributing to this protective effect.

**Conclusions:** We developed an adhesion-based high-throughput screening platform capable of identifying adhesion modulators. We revealed and validated several pro-adhesive compounds and showed a protective effect for the top candidate dexamethasone in an ACM mouse model. This indicates the potential of adhesion-strengthening compounds as therapeutic strategy in ACM and lays the basis for detailed follow-up studies. Moreover, these results and methods can be translated to other diseases with defective desmosomal adhesion.

**Clinical Perspective:** *What is new?:*

- We developed a novel high-throughput *in vitro* screening platform to identify compounds that strengthen cell-cell adhesion under conditions of disrupted adhesion, which is a central pathogenetic step in Arrhythmogenic Cardiomyopathy (ACM).
- With aid of this screen, we identified and validated different compounds from an FDA-approved drug library, which restore disrupted cell-cell adhesion induced by ACM patient mutations including glucocorticoids such as dexamethasone.
- Dexamethasone rescued impaired right ventricular function in a novel inducible ACM mouse model and reverted altered AKT1/AMPKα1 signaling as well as changes in the phosphorylation state of components of the contractile and junctional apparatus.

*What are clinical implications?:*

- We established a high-throughput cell-cell adhesion screening method as a viable tool for identifying drugs to combat ACM and other diseases featuring impaired desmosomal adhesion.
- We identified the glucocorticoid dexamethasone as a promising compound to therapeutically approach ACM.
- This study highlights strengthening of disrupted cell-cell adhesion as a novel therapeutic strategy to treat ACM.

## Introduction

Arrhythmogenic Cardiomyopathy (ACM) is a genetic disorder characterized by ventricular arrhythmia, impaired cardiac function and altered electrical conduction. With a prevalence of 1:1’000 to 1:5’000, it is one of the major causes of sudden cardiac death in young adults.^1^ Structurally, ACM is characterized by progressive loss of cardiomyocytes and patch-like replacement by fibrotic tissue and ventricular dilatation.^1,2^ Because the underlying molecular and cellular mechanisms leading to arrhythmia and structural changes are insufficiently understood, current treatment options are only symptomatic ranging from restriction of physical exercise, pharmacological treatment with beta-adrenergic blockers and antiarrhythmic drugs, implantation of a cardioverter defibrillator, catheter ablation, and heart transplantation as the last option.^1^ This highlights the need for identification of more targeted treatment strategies and a better understanding of the underlying cellular and molecular mechanisms leading to ACM.

In the majority of ACM cases, mutations in components of the desmosomal cell-cell adhesion complex can be identified.^1^ In cardiomyocytes, desmosomes are located at the intercalated disc to promote mechanical coupling. This junctional complex consists of the cadherin-like transmembrane adhesion molecules desmoglein-2 (DSG2) and desmocollin-2 (DSC2), which are anchored to the desmin intermediate filament system via a set of desmosomal plaque proteins including plakoglobin (JUP), plakophilin-2 (PKP2) and desmoplakin (DSP). For each of the desmosomal genes, pathogenic and likely pathogenic variants were described within the entire coding sequence of the proteins.^3^ Dysfunctional intercellular adhesion was shown to be a crucial initial event in ACM pathogenesis and the development of the disease phenotype in an ACM mouse model bearing a targeted knock-in mutation to abrogate specifically the adhesive binding function of DSG2.^4^ In line with this, structural remodeling of the intercalated disc is a common feature in ACM patients.^5–8^

From this central relevance of compromised adhesion for the ACM pathomechanism, it can be deduced that strengthening of mechanical coupling can serve as a potential therapeutic strategy to avoid adverse downstream effects. Moreover, several studies highlight the feasibility of modulating cardiac cell-cell adhesion via modulation of signaling pathways such as β-adrenergic/cAMP/PKA or ERK/MAPK.^5–8^

Therefore, we here aim to identify and evaluate compounds with respect to their ability to alter the cell-cell adhesive state in the context of ACM. To achieve this, we established and validated a novel 2D high-throughput adhesion assay to quantify cell-cell adhesion *in vitro* and investigated an FDA-approved drug library with respect to the pro-adhesive function of the compounds. Based on the results of the screen and in conjunction with *in vitro* validation experiments, we identified a group of pro-adhesive compounds with the ability to strengthen disrupted intercellular adhesion under ACM-like conditions. Within the top pro-adhesive drugs, glucocorticoid receptor agonists (glucocorticoids) were enriched. *In vivo* evaluations in an ACM mouse model revealed a protective effect of the representative glucocorticoid dexamethasone through restoration of right ventricular systolic function.

This study proposes the repurposing of already approved drugs based on their ability to strengthen cell-cell adhesion as potential therapeutical approach in ACM and has further implications for other diseases presenting with defective cell-cell adhesion such as skin blistering diseases or inflammatory bowel diseases.^9–13^

## Methods

### Drugs and compounds

Following drugs were applied for validation experiments and incubated for 24 hours. Metformin hydrochloride (Metformin, HY-17471A, MedChemExpress (MCE), South Brunswick, USA), DL-alpha-tocopherol (HY-W020044, MCE), Diclofenac sodium (Diclofenac, HY-15037, MCE), Vildagliptin (HY-14291, MCE), Disodium succinate (Disod. succinate, HY-W015410, MCE), Dexamethasone disodium phosphate (Dexamethasone, HY-B1829A, MCE), all dissolved in H_2_O. Adefovir dipivoxil (Adefovir, HY-B0255, MCE), Siponimod (HY-12355, MCE), Cobimetinib hemifumarate (Cobimetinib, HY-13064A, MCE), Nedocromil (HY-13448, MCE), Medroxyprogesterone (HY-B0648, MCE), Chlorothiazide (HY-B0224, MCE), Sunitinib (HY-10255A, MCE), Rosiglitazone (HY-17386, MCE), Mirtazapine (HY-B0352, MCE), Miconazole nitrate (Miconazole, HY-B0454A, MCE), Dexamethasone acetate (HY-14648A, MCE), Tolfenamic acid (HY-B0335, MCE), Torsemide (HY-B0247, MCE), Ruxolitinib phosphate (Ruxolitinib, HY50858, MCE) were dissolved in dimethyl sulfoxide (DMSO, D2438 Sigma-Adrich (Merck), Munich, Germany). Drugs were applied at 1 nmol/l (chlorothiazide), 100 nmol/l (mirtazapine), 1 µmol/l (dexamethasone disodium phosphate) or 10 mM (all other drugs) concentration. The combination of the adenylyl cyclase activator forskolin and phosphodiesterase-4 inhibitor rolipram (F/R, Sigma-Aldrich) was applied at concentrations of 5 µmol/l and 10 µmol/l, respectively and were incubated for 60 min.

### Cell lines and cultivation

The human epithelial cell line CaCo2 was kindly provided by Nicolas Schlegel (Department of General, Visceral, Vascular and Pediatric Surgery, University Hospital Würzburg, Würzburg, Germany). Generation of a CaCo2 line expressing DSG2-W2A via a lentiviral construct was described in ^4^. All CaCo2 cell lines were maintained in Dulbecco’s Modified Eagle Medium (DMEM, D6546, Sigma-Aldrich) supplemented with 10 % fetal bovine serum (S0615, Merck, Darmstadt, Germany), 100 U/mL penicillin (VWR International, Radnor, USA) and 100 µg/mL streptomycin sulfate (VWR) and 2 mmol/l L-glutamine (Sigma-Aldrich) at 37 °C, 5 % CO_2_ and full humidity. The culture medium was changed every three days. For experiments, cells were seeded on TC-treated plastic cell culture plates, grown to confluency and cultivated for at least seven days.

The pancreatic ductal adenocarcinoma cell line AsPC-1 was cultured in RPMI 1640 Medium (1-41F01-I, Bioconcept, Allschwil, Switzerland) supplemented with 10% fetal bovine serum (S0615, Merck), 2 mmol/l stable glutamine (5-10k50-H, Bioconcept), 50 U/mL penicillin (VWR) and 50 µg/mL streptomycin sulfate (VWR). The culture medium was changed every three days. DSG2 KO clones were generated using the CRISPR/Cas9 technique as described in ^9^.

All cells were quarterly checked for mycoplasma contaminations using PCR and were proven negative. Cells were routinely authenticated by Short Tandem Repeat profiling.

### Plasmid generation, cloning and mutagenesis

#### Cloning of existing mutant constructs

Amplification was performed using the Takara Emerald PCR kit (RR310A, Takara Bio, Kusatsu, Japan) with respective PCR primers indicated below. Primers were designed and synthesized by Microsynth (Microsynth, Balgach, Switzerland). PCR was performed according to the manufacturer’s protocol. The PCR product was purified with Nucleospin gel and PCR clean up kit (740609, Macherey-Nagel, Düren, Germany). The PCR product and the pLenti-C-mGFP plasmid (PS100071, OriGene Technologies, Rockville, USA) were digested with respective enzymes overnight at 37 °C. The digested product was loaded on a 1% agarose gel and purified from the gel Nucleospin gel and PCR clean up kit. The insert was ligated into the vector with a ratio of 3:1 overnight at 16°C using T4 Ligase (300361, Bioconcept) and bacterial transformation was performed in DH5-α strains. The integrity of the plasmid, including the presence of the wanted mutation was verified by restriction and Sanger sequencing (Microsynth).

DSG2 constructs: Amplification was performed from full length human DSG2-WT, DSG2-A517V and DSG2-G812S in the pEGFP-N1 vector (kindly provided by Katja Gehmlich, University of Birmingham, ^14^) with PCR primers including an AscI and NotI restriction site and Kozak sequence. The PCR product and the pLenti-C-mGFP plasmid were digested with AscI (R0558L, New England Biolabs (NEB), Ipswich, USA) and NotI (300347, NEB). Generation of the murine DSG2-W2A-C-mGFP construct was described in ^4^.

DSC2 constructs: Amplification was performed from full length human DSC2-WT, DSC2-R203C and DSC2-T275M in the pEGFP-N1 vector (kindly provided by Katja Gehmlich, University of Birmingham, ^15^) with PCR primers containing a BamHI and XhoI restriction site and Kozak sequence. The PCR product and the plasmid pLenti-C-mGFP were digested with BamHI (300325, NEB) and XhoI (300366, NEB).

PKP2a-WT construct: Amplification was performed from full length human PKP2a-WT in the pD2509-neonGreen-Pkp2a vector (kindly provided by Judith Fülle and Christoph Ballestrem, University of Manchester, ^16^) with the PCR primers containing an AscI and MluI restriction site and Kozak sequence. The PCR product and the plasmid pLenti-C-mGFP were digested with AscI (R0558L, NEB) and MluI (R3198S, NEB). To prevent recircularization of the digested DNA fragment, the PCR products were treated with phosphatase alcaline and T4 Polynucleotide kinase (M0201S, NEB) according to the manufacturer’s protocol.

JUP-WT construct: Amplification was performed from full length human JUP-WT in the 330-JUP-Myc vector (32228, Addgene, Watertown, MA, USA) with the PCR primers including a AscI and NotI restriction site and Kozak sequence. The PCR product and the plasmid pLenti-C-mGFP were digested with AscI (R0558L, NEB) and NotI (300347, NEB).

DSP-WT construct: Generation of the pLenti-hDSP-WT-C-mGFP construct was described in ^17^.

#### Site directed mutagenesis

For ACM mutation plasmids not available from collaboration partners, the Q5 site directed mutagenesis kit (E0554S, NEB) was used and mutagenesis was performed as indicated by the manufacturer. 5 ng of template plasmid were used. The KLD reaction was incubated 45 min at room temperature. Table 1 indicates the template plasmids, the primer sequences, annealing temperature and extension time used for mutagenesis. The integrity of the clones and the presence of the mutation was verified by enzymatic restriction and Sanger sequencing.

**Table 1.**
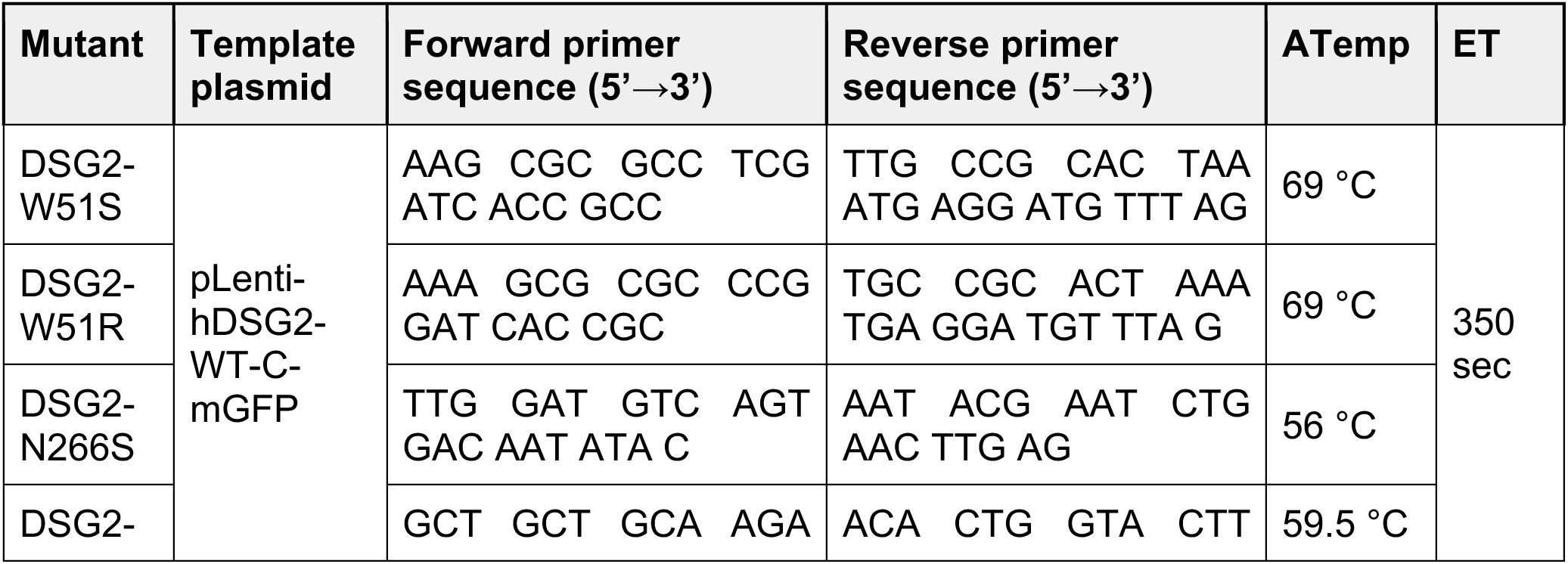
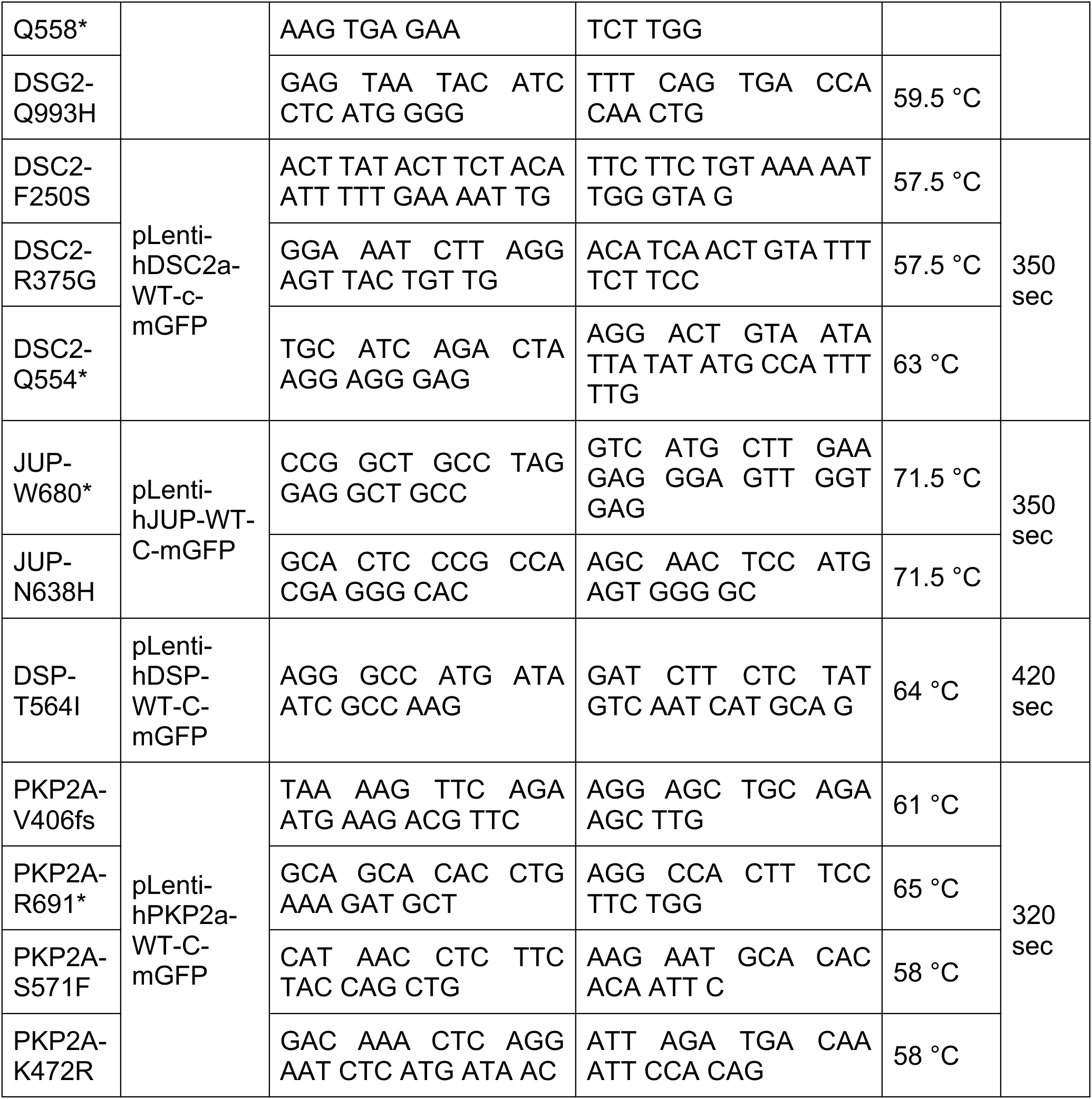
Annealing temperature (ATemp), Extension time (ET).

### Lentivirus generation and transduction

Lentiviral particles were generated according to standard procedures. In brief, HEK293T cells were co-transfected with the packaging vector psPAX2 (#12259, Addgene), the envelope vector pMD2.G (#12260, Addgene) and the respective construct plasmid using polyethylenimine (PEI Prime linear polyethylenimine, #919012, Sigma-Aldrich). After 48 hours, the supernatant containing virus particles was collected and enriched using LentiConcentrator (TR30026, OriGene). Cells were transduced with the respective concentrated virus particles using 10 µg/mL polybrene (H9268, Sigma-Aldrich) according to the manufacturer’s instructions. After 24 hours, the medium was changed and cells cultivated for at least one week before starting with the respective experiments. Expression of the respective construct was confirmed via visual detection of fluorescent signal and quantitative real-time PCR (qPCR) amplification of the mGFP reporter transcript.

### Standard adhesion assay

Cells were treated as indicated and grown to confluency in 24-well plates. Cell monolayers were washed with Hank’s balanced salt solution (HBSS, H8264, Sigma-Aldrich) and incubated with dissociation buffer (dispase II 2.5 U/mL, D4693, Sigma-Aldrich; DNAseI 10 ng/µl, A3778, Applichem, Darmstadt, Germany; in HBSS) at 37 °C till detachment of the cell monolayer from the well bottom (30 minutes). After detachment, monolayers were mechanically stressed by defined pipetting using an electrical pipette (Eppendorf Explorer, L49475G, Eppendorf, Hamburg, Germany). The total number of resulting fragments per well was determined using a binocular stereo microscope (SZX2, Olympus, Tokyo, Japan). Fragments were counted if they were clearly visible at 1.25-fold magnification. The number of fragments is an indirect measure for intercellular cohesion.

### High-throughput adhesion assay

30’000 AsPC-1 DSG2 KO cells were seeded on 96-well plates (TC-treated, Greiner Bio-One, Kremsmünster, Austria) in 200 µl medium using a P300 multichannel pipet (Eppendorf) and cultivated as indicated in *Cell lines and cultivation* for 7 days until they reached confluency. Medium was changed four days after seeding and 24 hours prior to the assay including respective addition of compounds. For the adhesion assay, cells were washed twice with HBSS and incubated with 50 µl dissociation buffer per well at 37°C for 1 hour. After detachment, defined mechanical shear stress was applied by rotation on a plate shaker (Microplate shaker model 980131CH, 3 mm orbit, VWR) at 750 rpm for 3 min with subsequent fixation and staining of the fragments with fixation solution (2 % paraformaldehyde, 10131580, Fisher Scientific, Schwerte, Germany; 20 µg/ml propidium iodine PI, 25535-16-4, VWR; 0.1 % Tween 20, BP337, Fisher Scientific). 48 to 96 hours after fixation, entire plates were scanned with a 4x objective mounted on an Olympus IX83 with a motorized stage and PI fluorescence was acquired.

### Drug library screen

The described high-throughput adhesion assay was used to screen an FDA-approved drug library (HY-L022M, MCE) containing drugs pre-dissolved at 10 mmol/l in DMSO or H_2_O in a 96-well plate format. For each assay run, AsPC-1 DSG2 KO were seeded on eight 96-well plates and processed in parallel. In this setup, we screened one drug-containing 96-well plate in three different concentrations plus vehicle control (1 nmol/l, 100 nmol/l, 10 µmol/l), each in duplicates. The compounds containing library plate was thawed at room temperature and centrifuged before use. Medium containing the respectively diluted compounds was prepared in 96-well deep plates (96-well deep PP plate, TreffLab, Degersheim, Switzerland) with a final vehicle concentration of 0.1 % in all conditions. For incubation of the cells, old medium was aspirated and the medium containing the respective drug was added sticking to the original compound library plate design. After incubation for 24 hours at 37 °C, *high-throughput adhesion assay* was performed as described above.

### Analysis screen data

Semi-automated analysis of fragment numbers was performed with the QuPath software (QuPath, Version: 0.5.1, The University of Edinburgh, ^18^). Position of the respective wells in the acquired plate images was defined, saved in .geojson format and applied accordingly. Number of fragments were analyzed applying the “Fragments detection” script (Link to GitHub script will be provided in the final version) based on the classifier function. For final number of fragments, results from the pixel classifiers (1) pre-filter “numerical opening” and (2) pre-filter “numerical closing” were averaged. Revealed number of fragments were normalized to the mean value of the respective vehicle control conditions from the same run and well row (A-H). The average values were calculated per drug concentration applied in duplicate. Drugs were ordered based on the maximum effect within the three tested concentrations.

### STITCH analysis

Top 100 drugs revealed by the high-throughput adhesion assay were analyzed with respect to their molecular action via the STITCH analysis tool (http://stitch.embl.de)^19^, at a minimum required interaction score of 0.400, with no more than 10 interactors on the 1st shell including all available active interaction sources. The revealed interactive interaction network was exported as .svg file.

### Generation of inducible ACM mouse model

All mouse experiments were carried out according to the protocol approved by the Cantonal Veterinary Office of Basel-Stadt (License number 3070) or the UKE Hamburg (N 064/2024) and complied with the ARRIVE guidelines. All mice were housed under specific pathogen-free conditions with standard chow and bedding with 12 hours day/night cycle according to institutional guidelines. To generate mice developing an inducible ACM phenotype, B6.129*Dsg2*tm1Mdcb,^9,20^ (kindly provided by the Max-Delbrück-Centrum für Molekulare Medizin in der Helmholtz-Gemeinschaft, Arnd Heuser) which homozygously express *loxP* sites flanking exon 2 of *Dsg2* were crossed with B6.Cg-Tg(CAG-cre/Esr1*)5Amc/J (004682, The Jackson Laboratory, Bar Harbor, USA), which express a tamoxifen inducible Cre recombinase. For genotyping of the animals, DNA was extracted from biopsies in 25 mmol/l NaOH and 0.2 mmol/l EDTA at 98 °C for 1 hour and neutralized with 40 mmol/l Tris pH 5.5. PCR was performed using GoTaq G2 (M7845, Promega, Madison, WI, USA) according to manufacturer’s instructions with the primers 5’-CCAGAGGAAACAACCTGGAA-3’ (forward) and 5’-GCACAGGACTCAGGATTGGT-3’ (reverse), which span the floxed region of *Dsg2*, and 5’-GCTAACCATGTTCATGCCTTC-3’ (forward) and 5’-AGGCAAATTTTGGTGTACGG-3’ (reverse) to detect the CAG-Cre transgene. Animals without the Cre transgene served as control group. Respective genotype of experimental animals was confirmed by qPCR.

### *In vivo* drug treatment

Mouse experiments were approved by the Cantonal Veterinary Office of Basel-Stadt (license numbers 3070) or the UKE Hamburg (N 064/2024). Animals of both sexes were applied without bias. For treatments, control and ACM mice from the same litter and sex were paired and allocated randomly to the treatment groups. For Cre induction, tamoxifen (T5648, Sigma-Aldrich) was dissolved in corn oil and administered at 75 mg/kg body weight intraperitoneally (i.p.) for 5 consecutive days. Dexamethasone disodium phosphate (HY-B1829A, MCE) was dissolved in PBS and administered three times per week at a concentration of 5 mg/kg body weight i.p. for the indicated time. Respective amount of PBS was administered to the treatment control group. During treatment, physical status of mice was checked daily. ECG measurements, echocardiography and heart sample collection and analysis were performed as described in the respective sections.

### Echocardiography and electrocardiogram (ECG)

Transthoracic echocardiography was performed using the Vevo 2100 ultrasound system (VisualSonics, Toronto, Canada) equipped with a MS-550 linear-array probe working at a central frequency of 40 MHz. After the animals were anesthetized with 3.0 % (v/v) isoflurane carried by pure oxygen, they were placed at supine position on a pre-warmed imaging platform. Anesthesia was maintained by 1.5 % (v/v) isoflurane through a nose cone and the body temperature was controlled at 37 °C by a rectal thermocouple probe. Eye gel (Lacrinorm) was applied to prevent ocular dehydration. Hairs on the chest were removed by applying commercially available hair removal cream (Nair). Left ventricle (LV) geometry and function were evaluated using 2D guided M-mode at the mid-papillary muscle level from parasternal short-axis (SAX). LV anterior (LVAW) and posterior (LVPW) wall thickness and internal dimensions (LVID) were measured from the M-mode during maximum systole (s) and diastole (d). Values were averages of three cardiac cycles. Left ventricular ejection fraction (EF) was calculated from derived volumes (Vol), which are computed based on the Teichholz formula (LV Vol;d = (7.0 / (2.4 + LVID;d)) × LVID;d^3^, LV Vol;s = (7.0 / (2.4 + LVID;s)) × LVID;s^3^, EF % = 100 ×((LV Vol;d – LV Vol;s) / LV Vol;d)). Left ventricular mass (LV Mass) was calculated based on a corrected cube model (LV Mass = 1.053 × ((LVAW;d + LVID;d + LVPW;d)^3^ – LVID;d^3^) × 0.8). To assess right ventricle (RV) function, RV fractional area change (RV FAC) was measured in B-mode at the mid-papillary level in SAX view. Briefly, the RV areas at end-diastole (RV Area;d) and end-systole (RV Area;s) were measured and RV FAC was calculated as 100 × (RV Area;d – RV Area;s) / RV Area;d (in %). Values were averages of three cardiac cycles. Data was transferred to an offline computer and analyzed with Vevo 2100 software (version 1.6.0, VisualSonics) by an investigator blinded to the study groups. For ECG recordings, mice were anesthetized as mentioned above, placed at supine position on a pre-warmed imaging platform and attached to the PowerLab Data Acquisition System (ML870 Powerlab 8/30, ADInstruments, Sydney, Australia). Needle probes were inserted subcutaneously in the right upper, and both lower limbs for acquisition of lead II. ECG was recorded for 5 min. ECG data were recorded and analyzed using the LabChart Pro 8 software (ADInstruments) equipped with the ECG Analysis Module. All mouse data were analyzed by an investigator blinded to the study groups. Peak amplitudes and intervals were determined as averages of three curve averages from 100 subsequent QRS complexes. After final measurements, mice were euthanized via cervical dislocation under anesthesia and hearts were dissected.

### Murine heart sample collection

For heart dissection, mice were euthanized via i.p. pentobarbital overdose or via cervical dislocation under isoflurane anesthesia according to guidelines of the Cantonal Veterinary Offices and Universities. Hearts were removed by lateral thoracotomy and directly immersed in ice-cold HBSS supplemented with 20 mmol/l 2,3-Butanedione monoxime (BDM, B0753, Sigma-Aldrich) and 0.5 mg/mL heparin (11483277, Fisher Scientific). Hearts were cut transversely at mid papillary level and fixed in 4 % paraformaldehyde in PBS for 16-24 hours with subsequent formalin-fixed paraffin embedding (FFPE) according to standard procedures or snap frozen for proteomics analysis.

### Sirius red fibrosis staining

Sirius red staining was performed according to standard procedures. In brief, FFPE sections were deparaffinized, washed in distilled water and stained with Coelestine blue iron-alum solution (15156.00500, Morphisto, Offenbach aM, Germany), followed by Mayeŕs hemalum solution (1.09249.0500, Sigma-Aldrich) and PicroSirius Red solution (13422.00500, Morphisto) for 30 min. Sections were washed in 70 %, 90 % and 100 % isopropanol and cleared in xylene and mounted in Roti-Histokit II (Carl Roth, Karlsruhe, Germany). Images of histological sections were acquired with a 20x objective mounted on a AxioScan Z1 slide scanner (Zeiss, Jena, Germany). The area of collagen was analyzed using the QuPath software. Areas of interest (i.e. interventricular septum, right and left ventricles) were annotated by a blinded experimenter, color channels were separated by deconvolution and total and fibrotic tissue area was determined by applying the “fibrosis analysis” script (Link to GitHub script will be provided in the final version).

### Flow cytometry viability assay

Cells were cultivated as described for the adhesion assays in both, the standard format (24-well plate) and the high-throughput format (96-well plate). Following the indicated treatment, cells were subjected to the respective adhesion assay according to the given protocol. Zombie red live/death staining dye (423109, Biolegend, San Diego, USA) was applied to assess cell viability at different steps during the adhesion assay: (1) prior to dispase detachment/shear stress, (2) dispase detachment without application of shear stress and (3) full protocol with dispase detachment and application of shear stress. Zombie red dye was applied on cells for 20-60 minutes. A respective unstained control was collected in parallel. As positive control, cells were cooked for 5 min at 100 °C. After staining, cells were dissociated by 0.06 % trypsin (T/3760/48, Fisher Scientific)/ 0.02 % EDTA (20309.296, VWR) and incubated for 20 min at 37 °C. After single cell suspension was obtained, cells were washed in FACS buffer (PBS + 2 % FCS (Fetal Calf Serum, 2-01F00-I, Bioconcept) + sodium azide (S2002, SigmaAldrich) and fixed in 4 % paraformaldehyde for 10 min. Cells were washed twice, resuspended in 100 µl of FACS buffer per sample. Flow cytometry data were acquired with a Cytoflex S (V4-B2-Y4-R3, C09766, Beckman Coulter, Brea, USA) and 5’000 events were recorded for each sample. To determine the fraction of viable cells, overall events were gated for cell-size particles using SSC-A vs. FSC-A followed by FSC-Width vs. FSC-A to remove multiplets and gate on single cells. From the singlet population, the live cell gating is based on Zombie Red staining, including an unstained control and positive (dead cells) control, as shown in Supp. Fig. 1A. Data were analyzed using the FlowJo software (version 10.9.0, Becton, Dickinson and Company, Ashland, USA).

### RNA isolation

Cells were cultivated according to the standard protocol until they reached confluency. Cells were washed with ice-cold PBS and lysed in TRI reagent (93289, Sigma-Aldrich). RNA was isolated via the Direct-zol RNA MiniPrep kit (R2050, Zymo research, Irvine, USA), including DNAse digestion according to the manufacturer’s protocol. RNA isolation from FFPE heart sections was performed with the RNeasy kit (73504, Qiagen, Hilden, Germany) according to manufacturer’s instructions.

### Quantitative real-time PCR (qPCR)

RNA was isolated as described in RNA isolation. Quantity and quality of RNA was determined by Nanodrop 1000 Spectrophotometer (Thermo Fisher Scientific, Waltham, USA). Up to 2 µg of isolated RNA was used for reverse transcription with (1) SuperScript III (Thermo Fisher Scientific) for RNA from cultivated cells or (2) iScript cDNA Synthesis kit (1708890, Bio-Rad, Hercules, USA) for RNA from FFPE sections. The obtained cDNA was diluted and quantitative real time PCR was performed with (1) StepOne Real time PCR Systems (Applied Biosystems, Thermo Fisher Scientific) with an annealing temperature of 55 °C, and a total 45 cycles or (2) BioRad Real-Time System CFX384 with an annealing temperature of 60 °C, and a total 40 cycles. (1) Power SYBR Green PCR Master Mix (Thermo Fisher Scientific) or (2) iTAG Universal SYBR Green Master Mix (Bio-Rad) and the respective forward and reverse primers at a dilution of 10 µmol/l were used. Primers are listed in table 2 below and were all validated by a standard curve of the respective sample type. Values were normalized to the Ct value of the average value of the indicated reference genes (^#^).

**Table 2.**
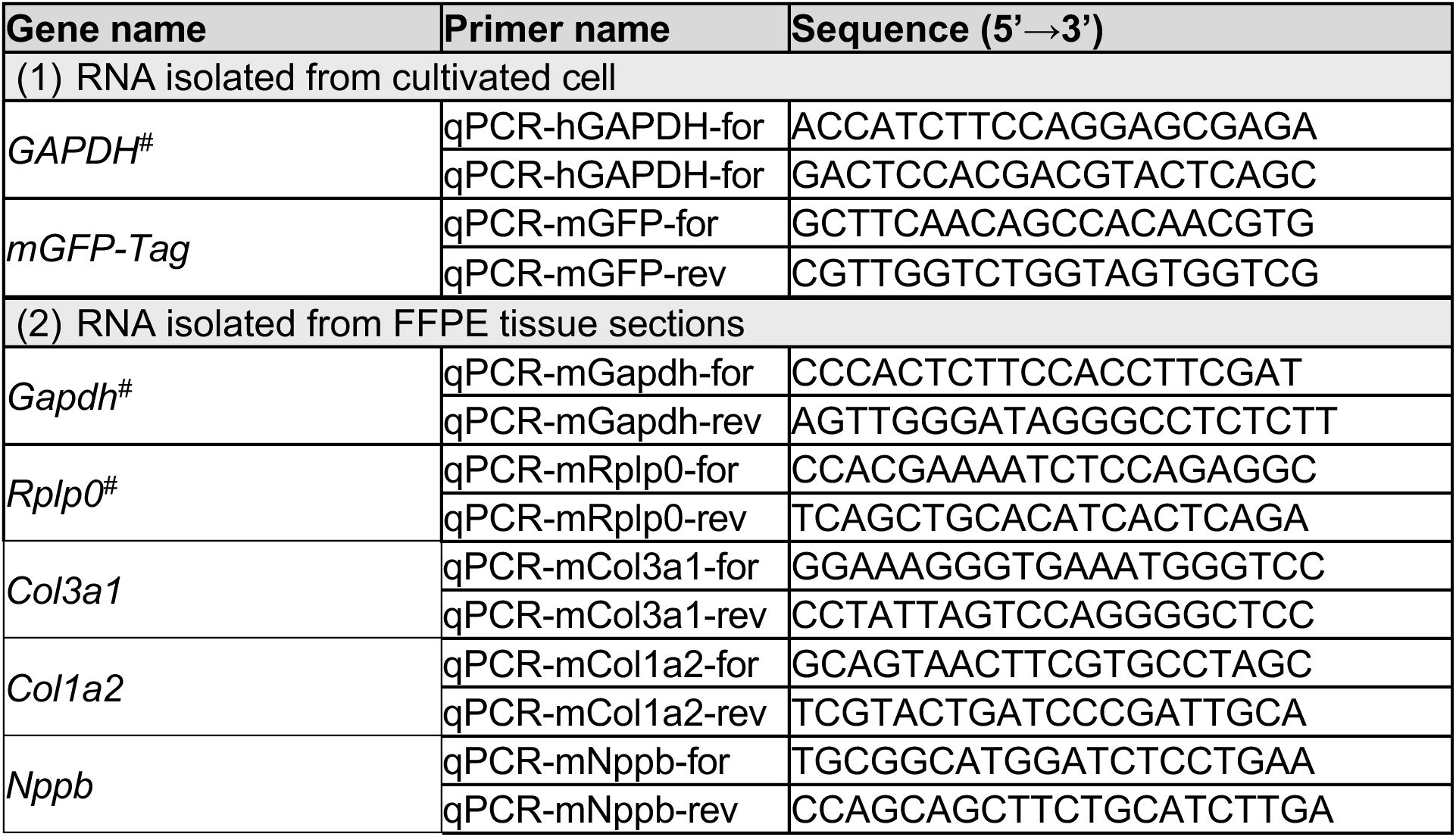

### Proteomics with PTM analysis

#### Sample preparation

Cardiac tissue was washed in ice-cold HBSS, frozen on dry ice and lysed in 2 mol/L Guanidinium-HCl (Gua-HCl), 0.1 mol/L ammonium bicarbonate, 5 mmol/L tris(2-carboxyethyl)phosphine (TCEP) and phosphatase inhibitors cocktail (Sigma-Aldrich, P5726 & P0044) by sonication (Bioruptor, 10 cycles, 30 seconds on/off, Diagenode, Belgium). Sonication was followed by the reduction for 10 min at 95 °C and then proteins were alkylated with 10 mmol/l chloroacetamide for 30 min at 37 °C. After diluting samples with 100 mmol/L ammonium bicarbonate buffer to a final Gua-HCL concentration of 0.5 mol/L, proteins were digested by incubation with sequencing-grade modified trypsin (1/50, w/w; Promega, Madison, Wisconsin) for 12 hours at 37 °C. After acidification using 5 % TFA, peptides were desalted using C18 reverse-phase spin columns (Macrospin, Harvard Apparatus) according to the manufacturer’s instructions, dried under vacuum and stored at –20 °C until further use. Peptide samples were enriched for phosphorylated peptides using Fe(III)-IMAC cartridges on an AssayMAP Bravo platform as previously described in ^21^. Both elution (of phosphorylated peptides) and flow-through (non-modified peptides) were collected and analyzed by mass spectrometry (MS).

#### Data acquisition and analysis of phosphorylated peptides

Phospho-enriched peptides were resuspended in 0.1 % aqueous formic acid and subjected to LC–MS/MS analysis using a Orbitrap Eclipse Mass Spectrometer fitted with an Ultimate 3000 nanoLC (both Thermo Fisher Scientific) and a custom-made column heater set to 60 °C. Peptides were resolved using a RP-HPLC column (75μm × 36cm) packed in-house with C18 resin (ReproSil-Pur C18–AQ, 1.9 μm resin; Dr. Maisch GmbH) at a flow rate of 0.3 μL/min. The following gradient was used for peptide separation: from 2 % B to 8 % B over 5 min, to 20 % B over 45 min to 25 % B over 15 min to 30 % B over 10 min to 35 % B over 7 min to 42 % B over 5 min to 50 % B over 3min to 95 % B over 2 min followed by 18 min at 95 % B. Buffer A was 0.1 % formic acid in water and buffer B was 80 % acetonitrile, 0.1 % formic acid in water. The mass spectrometer was operated in DDA mode with a cycle time of 3 seconds between master scans. Each master scan was acquired in the Orbitrap at a resolution of 120’000 FWHM (at 200 m/z) and a scan range from 375 to 1600 m/z. Maximum injection time was 50 ms and normalized AGC target was 250 %. Each MS scan was followed by MS2 scans of the most intense precursors in the Orbitrap at a resolution of 30’000 FWHM (at 200 m/z) with isolation width of the quadrupole set to 1.4 m/z. Maximum ion injection time was set to 54 ms (MS2) with an AGC target set “Standard”.

Only peptides with charge state 2 – 5 were included in the analysis. Monoisotopic precursor selection (MIPS) was set to Peptide, and the Intensity Threshold was set to 2.5e4. Peptides were fragmented by HCD (Higher-energy collisional dissociation) with collision energy set to 30 %, and one microscan was acquired for each spectrum. The dynamic exclusion duration was set to 30 sec. The acquired raw files were imported into the Progenesis QI software (v2.0, Nonlinear Dynamics Limited), which was used to extract peptide precursor ion intensities across all samples applying the default parameters. The generated mgf file was searched using MASCOT against a murine database (downloaded from Uniprot on 2022.02.22) and commonly observed contaminants using the following search criteria: full tryptic specificity was required (cleavage after lysine or arginine residues, unless followed by proline); 3 missed cleavages were allowed; carbamidomethylation (C) was set as fixed modification; oxidation (M) and phosphorylation (STY) were applied as variable modifications; mass tolerance of 10 ppm (precursor) and 0.02 Da (fragments). The database search results were filtered using the ion score to set the false discovery rate (FDR) to 1 % on the peptide and protein level, respectively, based on the number of reverse protein sequence hits in the datasets. Quantitative analysis results from label-free quantification were processed using the SafeQuant R package v.2.3.2. (https://github.com/eahrne/SafeQuant/ ^22^) to obtain peptide relative abundances. This analysis included global data normalization by equalizing the total peak/reporter areas across all LC-MS runs, data imputation using the knn algorithm, summation of peak areas per protein and LC-MS/MS run, followed by calculation of peptide abundance ratios. Only isoform specific peptide ion signals were considered for quantification. To meet additional assumptions (normality and homoscedasticity) underlying the use of linear regression models and t-tests, MS-intensity signals were transformed from the linear to the log-scale. The summarized peptide expression values were used for statistical testing of between condition differentially abundant peptides. Here, empirical Bayes moderated t-tests were applied, as implemented in the R/Bioconductor limma package (http://bioconductor.org/packages/release/bioc/html/limma.html). The resulting per protein and condition comparison p-values were adjusted for multiple testing using the Benjamini-Hochberg method.

#### Data acquisition and analysis of flow-through fraction

Flow-through peptides were resuspended in 0.1 % aqueous formic acid and subjected to LC–MS/MS analysis using the same instrument as above with the following exception: The separation gradient was: from 2 % B to 12 % B over 5 min, to 30 % B over 40 min to 50 % B over 15 min to 95 % B over 2 min followed by 18 min at 95 % B. Buffer A was 0.1 % formic acid in water and buffer B was 80 % acetonitrile, 0.1 % formic acid in water.

The mass spectrometer was operated in DIA mode with a cycle time of 3 seconds. MS1 scans were acquired in the Orbitrap in centroid mode at a resolution of 60’000 FWHM (at 200 m/z), a scan range from 350 to 1200 m/z, normalized AGC target set to 250 % and maximum ion injection time mode set to 50 msec. MS2 scans were acquired in the Orbitrap in centroid mode at a resolution of 15’000 FWHM (at 200 m/z), precursor mass range of 400 to 900, quadrupole isolation window of 12 m/z with 1 m/z window overlap, a defined first mass of 120 m/z, normalized AGC target set to 800 % and a maximum injection time of 22 ms. Peptides were fragmented by HCD (Higher-energy collisional dissociation) with collision energy set to 33 % and one microscan was acquired for each spectrum.

The acquired files were searched using the Spectronaut (Biognosys v18.1) directDIA workflow using standard settings. The search was done against a Mus Musculus database (downloaded from Uniprot on 2022.02.22) and commonly observed contaminants. Quantitative fragment ion data (F.Area) was exported from Spectronaut and analyzed using the MSstats R package v.4.8.6.^23^. Data was normalised using the default normalisation option “equalizedMedians”, imputed using “AFT model-based imputation” and p-values and q-values for pairwise comparisons were calculated using the default settings of MSstats package.

### Data analysis proteomics

All analyses in R were performed using R version 4.3.2.

#### Proteomics (MSstats)

We processed the Spectronaut output using the MSstats R package (version 4.10.1). Parameters were set to remove proteins with only 1 feature, use “0” to denote censored missing values and use the top 1’000 features. Other parameters set to default. When a protein received multiple annotations (multiple Uniprot IDs), only the 1st one was kept. Quality control was performed, normalization effects assessed, and model assumptions checked as per the manual.

#### Phospho-proteomics

Oxidations were not considered, only phosphorylation, and as such were filtered out following the PCA and volcano plot analyses. For some proteins, one or more PTM’s were detected in the phospho-proteomics dataset even though the protein itself was not detected in the protein dataset. This is presumably due to the enrichment for phospho-proteins prior to the mass spectrometry step for these experiments.

#### Principal Component Analysis (PCA)

PCA analysis performed using the factoextra R package (version 1.0.7). For the proteomics data, proteins with missing values are omitted before running the algorithm. For the phospho-proteomics data, analysis was performed on unduplicated PTMs. Volcano plot: Volcano plots created using the EnhancedVolcano R package (version 1.20.0). Thresholds were placed at ≤ −1 and ≥ 1 for log_2_ fold change and 0.05 for the p-value (not adjusted for multiple testing). PTMs were not deduplicated for the phospho-proteomics data.

#### GO analysis

Gene ontology (GO) analysis for phospho-proteomic data calculated using the “enrichGO” function from the clusterProfilerR R package (version 4.10.1). Filtered for proteins with at least 1 PTM with log_2_ fold change of ≤ −1 or ≥ 1 in ACM Dexa mice compared to Control PBS mice, without deduplication of PTMs. Background was all the proteins found in the phosphoproteomic dataset. Functional enrichment analysis was performed for the Biological Process, Molecular Function, and Cellular Component categories. P-values were adjusted for multiple testing using the Benjamini and Hochberg step-up procedure, with FDR controlled at the 0.05 level. The resulting list of enriched GO terms was simplified using the “pairwise_termsim” function from the enrichplot R package (version 1.22.0) and “simplify” from clusterProfilerR.

#### PTM-SEA

For each peptide with one or more PTMs, per comparison, a score was calculated as in ^24^: peptide_score = −10 * log_10_(q-value) * sign(log_2_(foldChange)). Given that one peptide can have multiple PTMs and that one PTM for a given protein can occur on multiple peptides, the PTMs were de-duplicated to obtain unique monophosphorylated peptides as follows: when a monophosphorylated peptide version of a PTM exists, that one is used. If not, the version of the PTM on the peptide carrying the fewest PTMs was used. In this case, each PTM on that peptide receives the score of the total peptide for the de-duplicated PTM. Finally, when a PTM occurred on >2 peptides with equal number of PTM’s, we used the mean of the peptide scores for the deduplicated PTM. We then used Morpheus (http://software.broadinstitute.org/morpheus/) to cast the data into the correct format and used the R GUI version of PTM-SEA (version 10.1.0). The min.overlap parameter was set to 5, other parameters were kept to default. The mouse PTM signature set database used was “ptm.sig.db.all.uniprot.mouse.v2.0.0.gmt” (v2.0.0, 2022-10-25).

#### Ingenuity Pathway Analysis (IPA)

For both kinases of interest (AKT1 and AMPKα1), the proteins from the PTMsigDB kinase signature sets which overlapped with our PTM dataset were used as inputs for Ingenuity Pathway Analysis (version 24.0.1)^25^ to determine pathway connections. PTM’s were deduplicated as before, but log_2_ fold changes between ACM Dexa and ACM PBS were used instead of peptide scores. These were loaded into IPA and analyzed using a cutoff of 1.6. Using the Path Designer tool, the kinase pathway proteins were individually added. Fold change values were overlaid onto the network. Finally, connections were determined using the ‘Connect’ tool with the following settings: Direct interactions, Mammal species, other settings left to default. Proteins that were not connected to anything in the pathway were not plotted for the figure.

### Statistics and data Compilation

Figures were compiled with Adobe Photoshop CC 2025 and Adobe Illustrator CC 2025. Statistical computations were performed with Prism 10 (GraphPad Software, Boston, USA). For comparison of two or multiple groups, distribution of data was analyzed by a Shapiro-Wilk normality test, and group variances were analyzed by an F-test or a Brown-Forsythe test, respectively. According to the results of these tests, a parametric or non-parametric test with or without Welch’s correction for unequal variances was applied. The statistical test used to compare the respective data sets is described in the corresponding figure legend. Statistical significance was assumed at p < 0.05 and is indicated with *. Unless otherwise stated, data are presented as box from the 25th to 75th percentile with median indicated as line and whiskers from the data minimum to maximum with each dot representing the mean value of all technical replicates for the respective biological replicate. Each animal or independent seeding of cells was taken as biological replicate, or as indicated in the legend.

### Data Availability

Phospho-proteomics data were deposited in the public database GEO/ProteomeXchange Consortium via the PRIDE partner repository (accession number will be provided in the final version). QuPath scripts are available on GitHub as indicated. Additional data pertaining to the current article are available from the corresponding author upon request.

## Results

### Mutations derived from ACM patients induce loss of cell-cell adhesion

Impaired desmosomal adhesion is sufficient to induce the characteristic ACM disease features *in vivo* as revealed by a recently established knock-in mouse model bearing a point mutation abrogating the molecular binding mechanism of DSG2 (mouse p.W56A; mature protein: DSG2-W2A)^4^. To gain more insight into the clinical relevance of defective adhesion in the context of ACM, we evaluated the impact of patient-derived variants in desmosomal genes on the adhesive function. We expressed a set of desmosomal mGFP-fusion proteins bearing ACM patient-derived mutations and performed a cell-cell adhesion assay *in vitro*. In this assay, a confluent cell monolayer is detached via dispase II-based enzymatic digestion and subjected to defined mechanical stress resulting in fragmentation of the floating monolayer. The number of fragments correlates with the overall cell-cell adhesion strength in an inverse manner (Fig. 1A).^5^ We selected a set of genetic variations reported from ACM patients, which are classified as pathogenic, likely pathogenic or variant of unknown significance (Supp. Table 1) according to the ClinVar data base and which are located in different regions of the transmembrane adhesion molecules DSG2 and DSC2, and the desmosomal plaque proteins JUP, DSP and PKP2 (Fig. 1B). We cloned respective human mutant and wildtype (WT) mGFP-fusion proteins and lentivirally overexpressed the constructs in the human CaCo2 cell line with confirmation of expression by qPCR. As positive control, we included cells expressing murine DSG2-W2A, as this mutation was shown to specifically disrupt desmosomal adhesion and to induce the ACM phenotype in a mouse model.^4^ Importantly, expression of patient mutations affecting the same amino acid in the human protein (DSG2-W51S, DSG2-W51R) induced a similar adhesive defect with increased monolayer fragmentation after mechanical stress compared to expression of the respective WT protein (Fig. 1C, D). Moreover, expression of several other patient-derived mutations in different regions for each of the tested desmosomal molecules led to a loss of cell-cell adhesion, whereas some mutations did not alter cell-cell adhesion regardless of their classification. Whether this lack of effect was due to methodological limitations of the assay or to differences in mechanisms, in that other initial pathological factors may play a role in addition to defective adhesion, is not clear. Nevertheless, these data show that certain patient-derived ACM mutations in different proteins cause defective cell-cell adhesion when overexpressed in the 2D cell setting.

**Figure 1.**
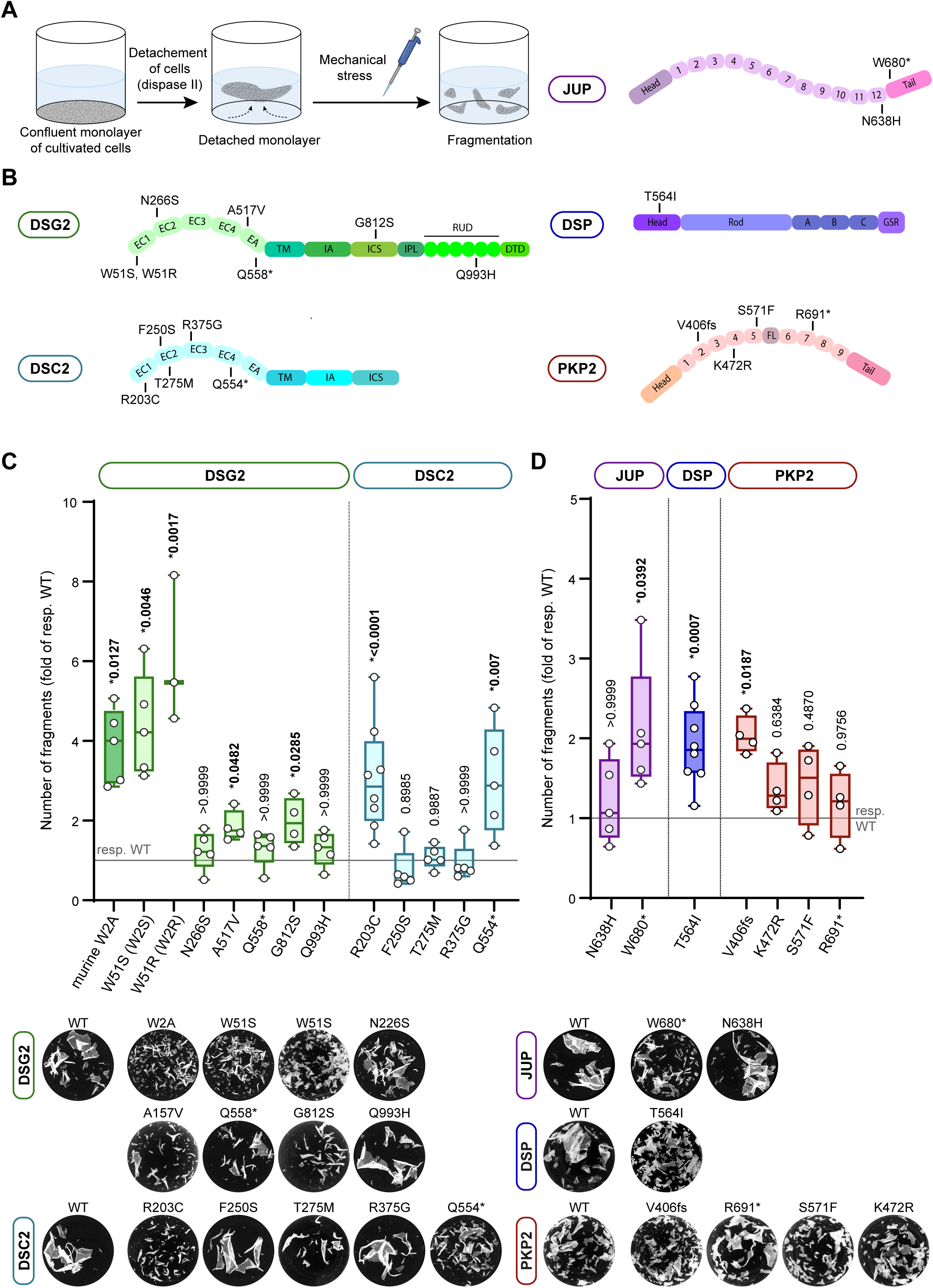
ACM-patient derived mutations disrupt cell-cell adhesion. A) Schematic displaying the steps of the cell-cell adhesion assay to determine intercellular adhesion in 2D cell culture. In this assay, a confluent CaCo2 cell monolayer is detached by enzymatic digestion via dispase II and defined mechanical stress is applied via electrical pipetting followed by evaluation of monolayer fragmentation via counting of the fragment number. B) Schematic depicting the position of the selected ACM patient mutations applied in this study. Extracellular domain (EC), extracellular anchor (EA), transmembrane domain (TM), intracellular anchor domain (IA), intracellular catenin-binding site (ICS), intracellular proline-rich linker (IPL), repeated unit domain (RUD), DSG-terminal domain (DTD), ARM repeats (1, 2, …), flexible insert (FL), central rod domain (Rod), phosphotransferase system regulation domain (A, B, C), glycine-serine-arginine rich (GSR) according to ^44^. C, D) Standard adhesion assay of CaCo2 cells overexpressing the indicated patient-derived ACM mutations in the transmembrane adhesion molecules desmoglein-2 (DSG2) and desmocollin-2 (DSC2) (C), and in the desmosomal plaque proteins plakoglobin (JUP), plakophilin-2 (PKP2a) and desmoplakin (DSP) (D), with representative images of respective wells after application of mechanical stress shown below. Number of fragments were normalized to the corresponding control condition (overexpression of the wildtype (WT) gene). Each dot represents a biological replicate (independent cell seeding) as mean of 3-4 technical replicates. Kruskal-Wallis test with Dunn’s correction or ordinary one-way ANOVA with Dunnett’s post-hoc test for multiple tests or one-sample t-test for a single mutation per gene for comparisons versus respective WT control.

### Establishment of a high-throughput adhesion assay

Together with the data from the DSG2-W2A mouse model^4^, these results indicate that dysfunctional desmosomal adhesion is a common, initial step during the development of ACM independent from the mutation-affected gene. To aid with the identification of new therapeutic strategies, we therefore aimed to detect compounds strengthening intercellular adhesion under ACM-like conditions with compromised mechanical coupling. To achieve this in a high-throughput format with the ability to evaluate compound libraries, we designed a screening platform based on the cell-cell adhesion assay described in Fig. 1A. In contrast to this standard assay employing 24-well plates, well-by-well application of shear stress and manual counting of the fragments, cells are cultivated in a 96-well plate format, detached with dispase, and defined mechanical stress is subsequently applied to up to 8 plates in parallel via orbital rotation on a multi-plate shaker (Fig. 2A). Resulting fragments are then fixed and stained with propidium iodide for low-background imaging. The number of fragments is determined in a semi-automated detection pipeline based on imaging of the fluorescence staining, and a fragment detection script implemented in the QuPath software^18^. Due to a lack of suitable cardiomyocyte cell lines, we implemented this workflow with the AsPC-1 epithelial cell line, which exhibits a desmosomal composition similar to cardiac myocytes. To mimic defective adhesion in the context of ACM, we employed a CRISPR/Cas9 cell line deficient for DSG2, the major desmosomal adhesion molecule in the heart (DSG2 KO).^9^ Importantly, DSG2 deficiency was shown to induce an ACM phenotype *in vivo*.^26^ In initial feasibility experiments, we seeded DSG2 KO and respective WT control cells on 96-well plates and performed the workflow as described. As expected from data of standard adhesion assays^9^, DSG2 KO cells showed consistently high fragmentation following mechanical stress (Fig. 2B). Importantly, we observed homogeneous fragmentation throughout the wells of entire plates showing the technical feasibility and stability of the set-up. We applied WT and DSG2 KO cells to identify the optimal cell seeding density and shaking conditions (frequence and time) resulting in the highest fragmentation ratio DSG2 KO vs. WT. This was followed by determination of the fixation, staining, and image acquisition protocol. Results of the automated image analysis were compared to manual fragment detection to identify the settings and parameters (signal intensity, threshold, detection algorithm) revealing comparable numbers. As reduced cell viability can affect cell adhesion and the results of the assay, we confirmed for the identified conditions that the necessary handling steps during the experiments do not cause increased cell death (Supp. Fig. 1). After establishment of the experimental workflow, we validated the capacity of the high-throughput assay to detect pro-adhesive effects induced by chemicals in general and to a similar extent as the standard assay. Therefore, we treated DSG2 KO cells with forskolin and rolipram (F/R) to increase cAMP levels, as this was shown to robustly reduce cell fragmentation in different cell types.^5,27^ In both assays, F/R treatment led to significantly reduced monolayer fragmentation with a comparable ratio of the mean number of fragments DMSO vs. F/R (Fig. 2C-E). This confirms that the established high-throughput assay can detect strengthening of cell-cell adhesion in DSG2 KO cells comparable to the standard assay and is thus suitable as a screening platform.

**Figure 2.**
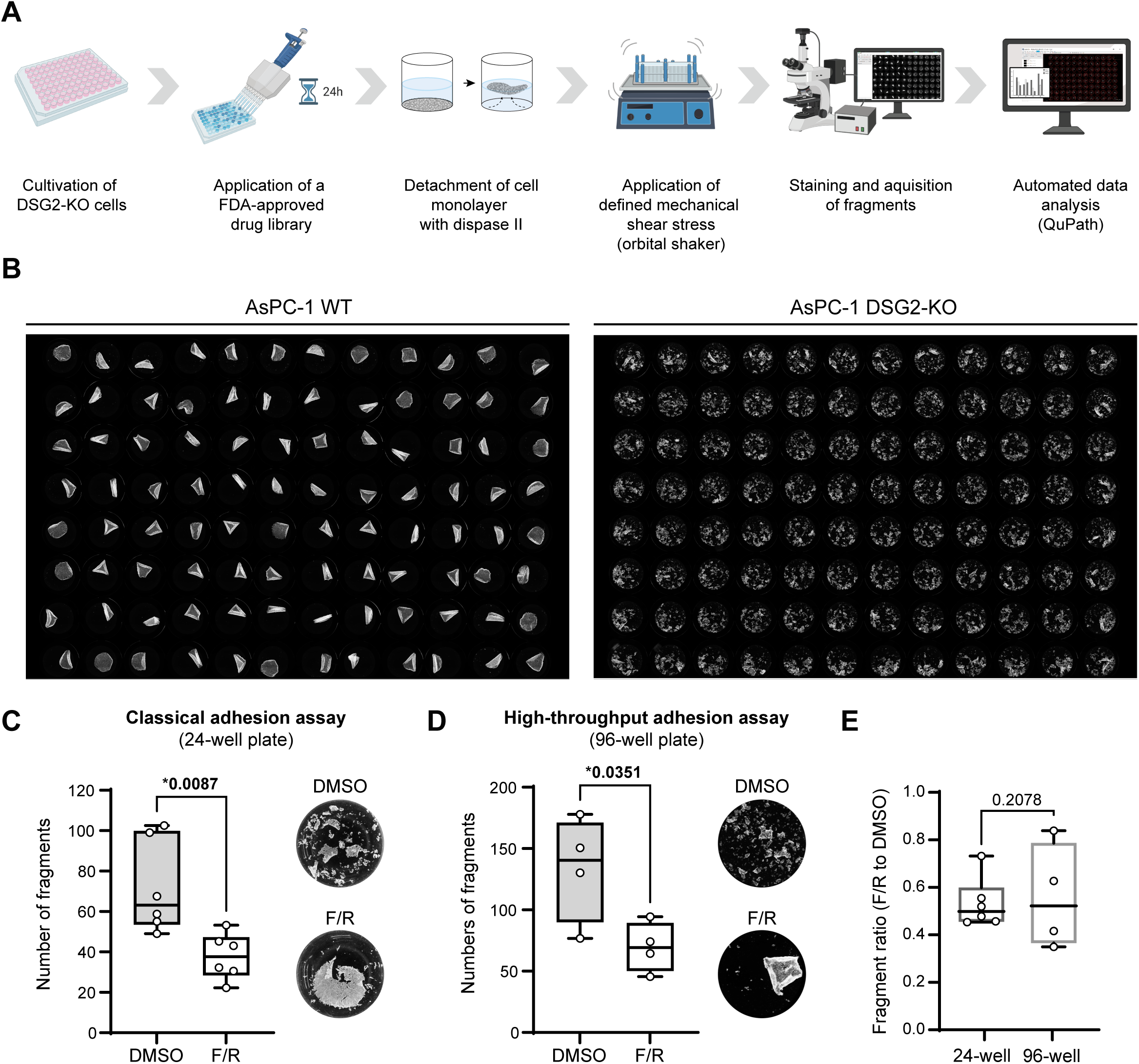
Establishment and validation of a high-throughput cell-cell adhesion platform. A) Schematic of the high-throughput cell adhesion assay platform. To screen a high number of agents, the standard cell-cell adhesion assay was adapted to a 96-well plate format with the ability to apply a compound library, including the establishment of a semi-automated pipeline for application of mechanical stress, automated image acquisition and analysis. B) Representative images of wildtype (WT) control AsPC-1 cells and DSG2 KO AsPC-1 cells seeded on 96-well plates and employed for initial feasibility experiments. C, D, E) Comparison of the standard adhesion assays in 24-well plate format to the high-throughput adhesion assay in 96-well plate format. Adrenergic stimulation by application of forskolin and rolipram (F/R) or DMSO vehicle control served as reference condition. Representative images of wells after shear stress are shown on the right. Each dot represents a biological replicate (independent cell seeding) as mean of 4 (for C), or 40 (for D) technical replicates, unpaired t-test. E) Ratio of fragment counts of F/R vs. DMSO derived from C, and D. Two-way ANOVA, with treatment and plate type as main factors and an interaction effect between the two.

### Screening of an FDA-approved drug library

In the next step, we employed this high-throughput adhesion assay platform to screen a compound library to identify drugs with the ability to increase cell-cell adhesion under the condition of disrupted desmosomal adhesion by DSG2 deficiency. Based on the idea of drug repurposing and fast transferability to the clinical context, we selected an FDA-approved drug library (MedChemExpress) containing 1’842 compounds with representatives from the majority of clinically used drug families and categories. To cover a large range of effective concentrations, we tested the drugs at 1 nmol/l, 100 nmol/l, 10 μmol/l, and the respective solvent (DMSO or water) as control with incubation for 24 hours prior to monolayer detachment. To correct for plate-to-plate variability, each condition was applied in duplicate on two different plates within a batch of 8 plates, all processed in parallel. The resulting mean fragment number per well was normalized to the respective mean control value from the same run. For downstream analysis, drugs were ranked according to their effect on cell-cell adhesiveness (-log_2_ fold change of fragment number) in DSG2 KO cells from high to low based on the maximum mean value revealed from the three tested concentrations per drug (Fig. 3A, Supp Table 2). From the tested conditions, 651 drugs induced a pro-adhesive effect with reduction of fragments by more than 25% for at least one concentration. In contrast, 354 compounds reduced cell-cell adhesion further under conditions of already compromised cohesion with an increase in the number of fragments of more than 25% in at least one of the concentrations.

**Figure 3.**
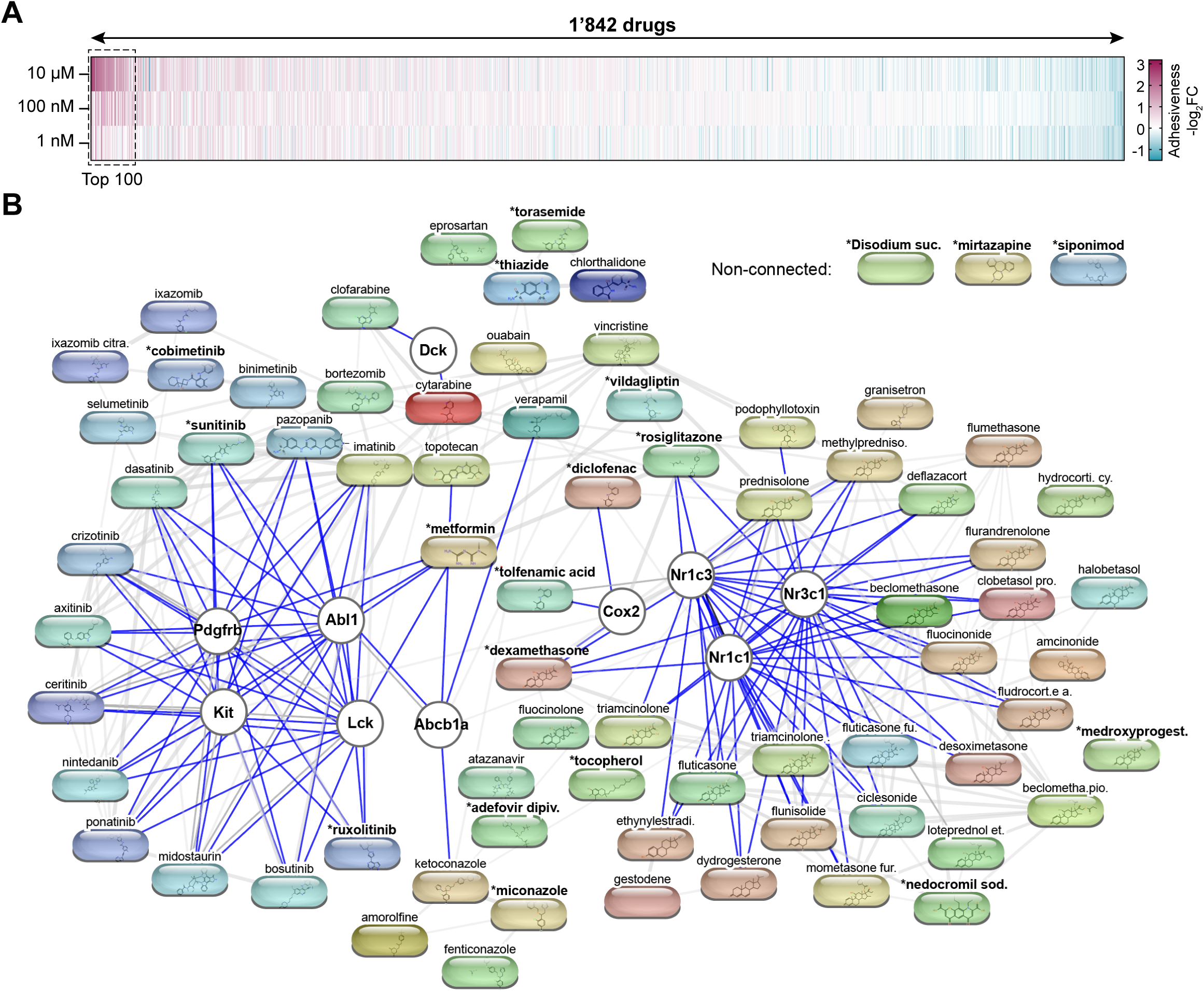
Drug library screen to identify compounds modulating cell-cell adhesion under ACM-like conditions. A) Heat map indicating the result of the high-throughput adhesion screen performed in DSG2 KO AsPC-1 cells. An FDA-approved drug library containing 1‘842 drugs was tested in three different concentrations (10 µmol/l, 100 nmol/l, 1 nmol/l) and normalized to respective vehicle control treatment. Data are displayed as adhesiveness defined as -log_2_ fold change of the revealed number of fragments normalized to the respective vehicle control. Drugs are ordered from left to right by decreasing maximal adhesiveness value among the tested concentrations. B) STITCH analysis of the top 100 compounds as indicated in (A) to identify common drug targets and interactions. Blue lines indicate direct binding of the drug to the respective protein. Compounds in bold marked with * were selected for validation experiments in Fig. 4.

As we intended to identify compounds which strengthen adhesion in the context of ACM, we focused on the top 100 pro-adhesive compounds for further analysis and aimed to identify possible overlapping downstream targets correlating with the pro-adhesive effect. Therefore, we performed a STITCH analysis of the top 100 drugs (Fig. 3B),^19^ which revealed two compound clusters sharing the same or related downstream targets with (1) a cluster of kinase modulators interacting with receptor tyrosine kinases (ABL1, LCK, KIT, PDGFRB) with high prevalence of tyrosine kinase inhibitors and (2) a cluster of modulators of nuclear receptors including the glucocorticoid receptor (NR3C1), peroxisome proliferator-activated receptor alpha and gamma (PPARα, PPARγ = NR1C1, NR1C3) with a dominance of receptor agonists. In line with this, 21 of the top 100 pro-adhesive drugs are known tyrosine kinase inhibitors or inhibitors of downstream effector kinases (JAK1/2, MEK1/2, ERK1/2), and 31 of the top pro-adhesive drugs are glucocorticoid receptor agonists and thus members of the glucocorticoid drug family. Together, this high-throughput adhesion drug screen revealed different drugs and drug classes increasing cell-cell adhesion under conditions with defective desmosomes with kinase inhibitors and glucocorticoids standing out.

### Validation of pro-adhesive compounds

Next, we evaluated the properties of the revealed top 100 pro-adhesive drugs more in detail with respect to severity of known side effects in patients, approved application format (exclusion of drugs with topical application only; oral administration is preferred over intravenous application), mode of action and molecular target. In conjunction with the results from the STITCH analysis and robustness of the pro-adhesive effect, we selected 20 compounds for validation experiments. To confirm the independence of the drug effect from (1) the type of cell line, (2) absence of DSG2, and (3) the high-throughput format, we conducted the adhesion assay in the standard format employing the CaCo2 cell line overexpressing DSG2-W2A (see Fig. 1C) and thus lacking DSG2 binding function with the full-length protein still being present (Fig. 4A). From the selected 20 compounds, 19 revealed an adhesion strengthening effect which was statistically significant versus control treatment for 7 drugs. From these compounds, we selected the 8 most pro-adhesive drugs based on median -log_2_FC and p-value and conducted a dilution series to determine the concentration range of the protective effect (Supp. Fig. 2). Dexamethasone was included as representative of the tested glucocorticoid receptor agonists (vs. dexamethasone acetate, medroxyprogesterone). For all drugs, a concentration-dependent effect was detectable, while dexamethasone and metformin showed a significant effect over 3 decimal powers of dilution indicating a broad therapeutical window with respect to the pro-adhesive effect.

**Figure 4.**
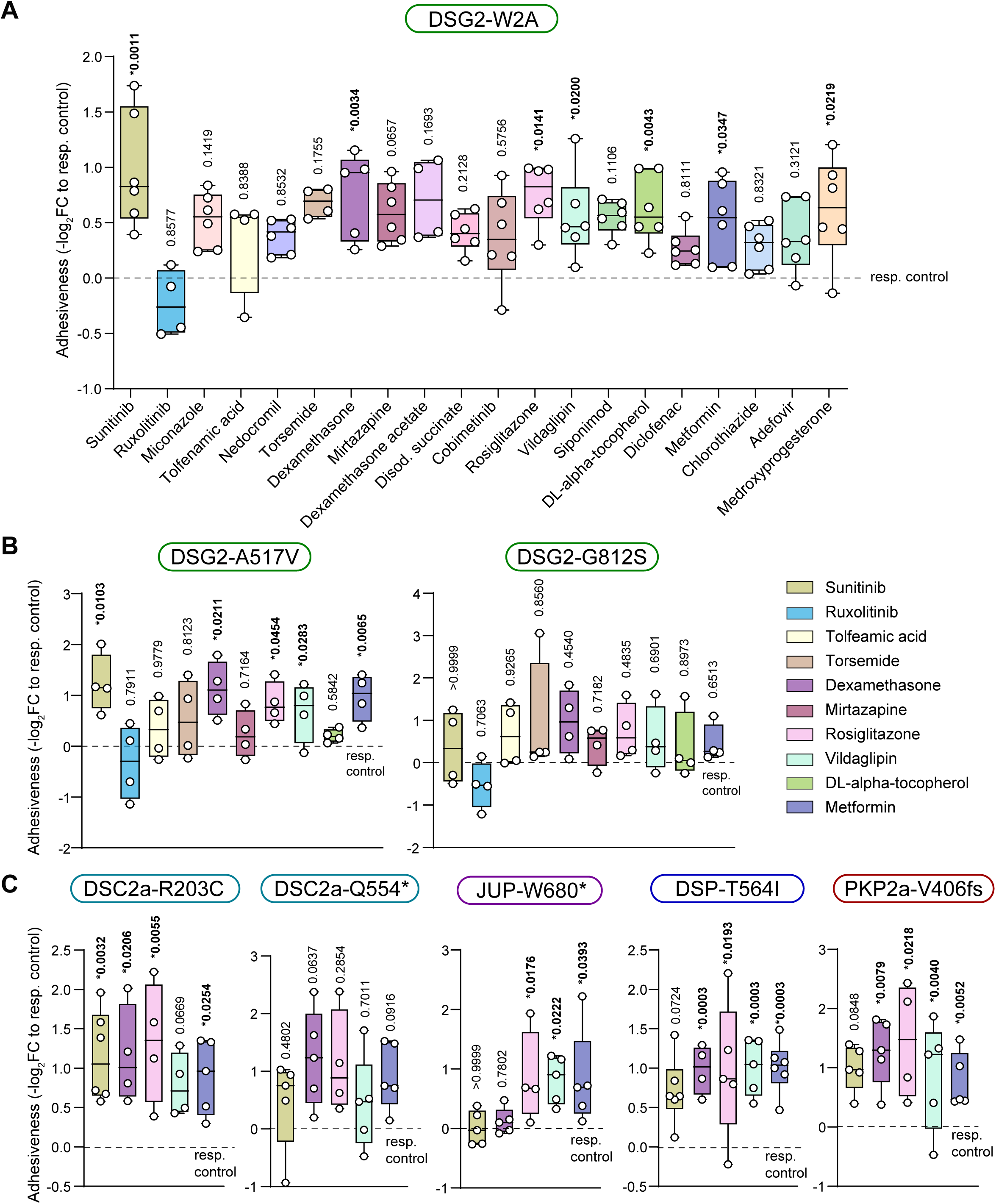
Validation experiments of pro-adhesive compounds revealed from an FDA-approved drug library. A) Validation of the top 20 selected drugs obtained from the screening experiment in Fig. 3. These were tested in standard adhesion assays in CaCo2 cell lines expressing the ACM mouse model DSG2-W2A mutation. Each dot represents a biological replicate (independent cell seeding) as mean of 4 technical replicates, results were normalized to the respected control treatment (dotted line), Kruskal-Wallis test with Dunn’s correction, ordinary one-way ANOVA with Dunnett’s correction or unpaired t-test vs. corresponding control condition was applied, respectively. B, C) Further evaluation of drugs selected from (A) in standard adhesion assays in CaCo2 cell lines expressing the indicated ACM patient mutations. Each dot represents a biological replicate (independent cell seeding) as mean of 2-4 technical replicates, results were normalized to the respected control treatment (dotted line), Kruskal-Wallis test with Dunn’s correction, ordinary one-way ANOVA with Dunnett’s correction or unpaired t test vs. corresponding control condition was applied, respectively.

In the screen and validation experiments, we employed cells bearing a DSG2 KO or DSG2-W2A mutation, respectively, to mimic defective cell adhesion as common pathogenic trunk in ACM. To analyze and confirm the potential of the selected drugs to strengthen adhesion also in the context of mutations derived from ACM patients, we employed the conditions shown in Fig. 1 C, D which exhibited disrupted adhesion. In addition to the 8 most pro-adhesive compounds, we included tolfenamic acid and ruxolitinib as representative drugs showing no effect or tending towards a negative effect in DSG2-W2A cells.

In cells expressing the ACM patient mutations DSG2-A517V or DSG2-G812S (Fig. 4B), the tyrosine kinase inhibitor sunitinib, the glucocorticoid dexamethasone, and the anti-diabetic drugs rosiglitazone, vildagliptin, and metformin significantly strengthened disrupted adhesion for DSG2-A517V cells, while this effect was not detectable for DSG2-G812S. For mirtazapine, torsemide, and DL-alpha tocopherol, no clear effect was detectable, and this was also the case for tolfenamic acid and ruxolitinib in both cell lines.

For the revealed top 5 pro-adhesive drugs, we extended our evaluations to ACM mutations in other desmosomal proteins (DSC2a-R203C, DSC2a-Q554*, JUP-W680*, DSP-T564I and PKP2-V406fs) (Fig. 4C) and found that dexamethasone, rosiglitazone, vildagliptin, and metformin reduced fragmentation for the majority of tested ACM mutations.

Thus, based on the several levels of selections, these 4 compounds showed the highest capacity to stabilize cell-cell adhesion under different ACM-like conditions and are the most interesting candidates for detailed follow-up analysis.

### Dexamethasone rescues cardiac function in an ACM mouse model

Based on the *in vitro* validation, we took one step further to the clinical translation and performed *in vivo* experiments to determine the impact of a pro-adhesive compound on ACM disease development and progression. Due to the strong enrichment of glucocorticoids within the top 100 pro-adhesive drugs, the well-known pharmacological properties, and the stable effect on different cell lines for a wide range of concentrations, we selected dexamethasone for a first pilot study in a novel ACM mouse model. The model is based on a CreER-mediated inducible deficiency of DSG2, which leads to a right ventricular dominant ACM phenotype over the course of 8 weeks. To achieve the maximum possible effect in this initial study, we started the application of dexamethasone or PBS control one week prior to induction of the disease and continued treatment via intraperitoneal injections three times a week for 8 weeks after induction (Fig. 5A). Development of the ACM phenotype was monitored via electrocardiogram (ECG) at baseline, 4, and 8 weeks after induction of the disease. As described earlier,^4^ progression to the disease was detectable by reduction of the QRS amplitude (Fig. 5B, C). Prior to termination of the animals 8 weeks after induction, cardiac function was evaluated via echocardiography with subsequent organ collection and histological analysis of cardiac fibrosis (Fig. 5B-E). In line with the ACM phenotype, mutant mice treated with PBS showed reduced systolic function of the right ventricle as assessed by the fractional area change (FAC) in conjunction with an elevated heart rate (Fig. 5D, Supp. Table 3). Interestingly, all parameters of left ventricular systolic function (ejection fraction, stroke volume, cardiac output) were not altered, despite detectable structural changes with cardiac fibrosis in both ventricles (Fig. 5B, E, Supp. Table 3). Under these conditions, treatment with dexamethasone improved right ventricular FAC and reverted QRS voltage reduction, while no clear effect on cardiac fibrosis was detectable (Fig. 5B-E). This finding is supported by mRNA analysis, which reveals that dexamethasone reduced the increased expression of *Nppb* (Natriuretic Peptide B) expression, a marker for increased cardiac wall tension and heart failure, in mutant hearts (Fig. 5F). In line with results from histology, elevated levels of collagens *Col1a2* and *Col3a1* were not altered (Fig. 5F). With respect to arrhythmia as a hallmark of ACM, 27.3 % of the mutant mice with control treatment exhibited premature ventricular contractions (PVCs) 8 weeks post disease induction, whereas this was slightly reduced to 23.1 % in dexamethasone treated mutant mice (Fig. 5G). However, the power of these data is limited due to the short ECG recording time of 5 minutes. In summary, these *in vivo* data highlight a protective effect of the pro-adhesive compound dexamethasone on certain functional characteristics of ACM, which suggests a potential relevance as therapeutic approach.

**Figure 5.**
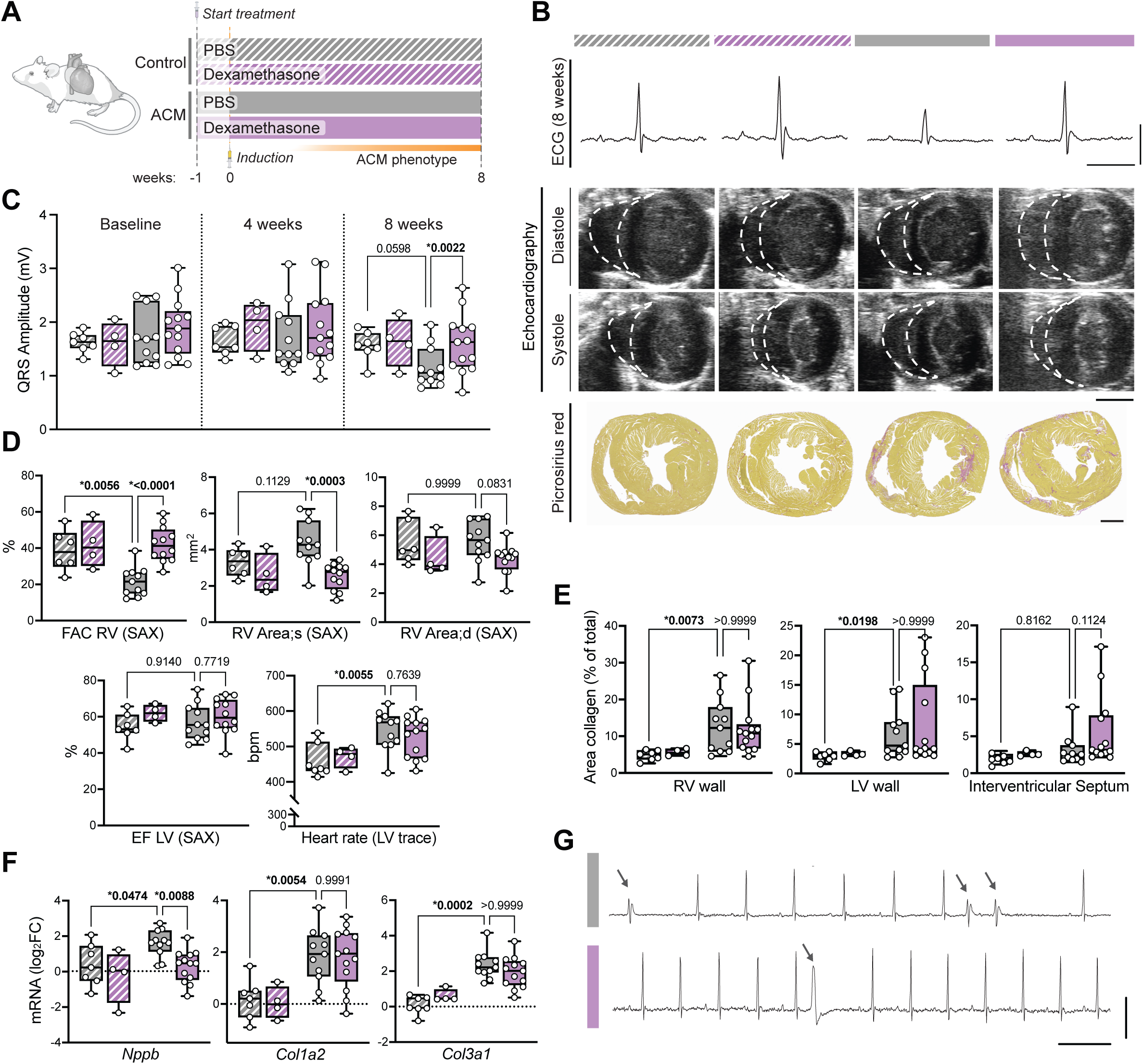
Dexamethasone treatment is protective in an ACM disease mouse model. A) Schematic depicting the experimental set-up for *in vivo* application of dexamethasone and PBS control to an inducible ACM mouse model. Treatment started one week prior to disease induction and continued for 8 weeks. Dexamethasone disodium phosphate was applied i.p. at 5 mg/kg body weight, three times a week. B) 8 weeks post-induction, the cardiac phenotype was evaluated by electrocardiogram (ECG), echocardiography and histological analysis. Color bars indicate the respective conditions according to (A). Representative QRS complexes from ECG analysis are shown in the top row, scale bar vertically = 1 mV, scale bar horizontally = 0.05 sec. The second and third row show representative images of echocardiography analysis in short axis view at mid papillary level in end systolic and diastolic position. White marks outline the right ventricle. Scale bar = 2 mm. In the last row, representative images of picrosirius red staining mark fibrotic areas. Scale bar = 1 mm. C) Analysis of the voltage amplitude of the QRS complex in ECG analysis at the indicated time points. Each dot represents the value of one animal, two-way RM ANOVA with Dunnett’s post hoc test, repeated measurements of same animal matched. D) Depicts selected parameters of echocardiography analysis acquired in short axis view (SAX) at mid-papillary level, including the fractional area change (FAC) and end-systolic (s) and diastolic (d) area as parameter for right ventricular (RV) output function, ejection fraction (EF) of the left ventricle (LV) and corresponding heart rate. See Supp. Table 3 for all analyzed parameters. Each dot represents the value of one animal, for RV Area;d, a Kruskal-Wallis test with Dunn’s post hoc test was performed, all other panels: ordinary one-way ANOVA with Sidak’s post hoc test. E) Analysis of cardiac fibrosis by determination of the area fraction of picrosirius red (= collagen) stained tissue versus total tissue area in the right ventricular (RV) wall, the left ventricular (LV) free wall and the interventricular septum. Each dot represents the value of one animal, Kruskal-Wallis test with Dunn’s post hoc test. F) Relative mRNA expression of the indicated genes in FFPE heart sections normalized to *Gapdh* and *Rplp0*. Each dot represents the value of one animal, *Col3a1*: Kruskal-Wallis test with Dunn’s post hoc test, all other panels: ordinary one-way ANOVA with Sidak’s post hoc test. G) Example ECG traces from ACM animals treated with PBS or dexamethasone presenting with premature ventricular contractions (PVCs) indicated by arrows. Scale bar vertically = 1 mV, scale bar horizontally = 0.1 sec.

### Altered phospho-proteomic profiles in response to Dexamethasone treatment

Finally, we aimed to determine the molecular mechanism underlying the protective effect of dexamethasone on cardiac function and cell-cell adhesion. As it was shown that post-translational modifications (PTMs) such as protein phosphorylation are not only key players for general heart functions (e.g. electromechanical coupling), but that they are also important modulators of cell-cell adhesion,^5,28,29^ we analyzed the total and phospho-proteome of ACM mouse ventricular tissue. To reduce the effect of secondary structural changes in heart morphology, we collected hearts of ACM mice treated with dexamethasone or PBS two weeks after induction of the disease and prior to development of cardiac fibrosis (Fig. 6A). As reference, control animals treated with PBS were included.

**Figure 6.**
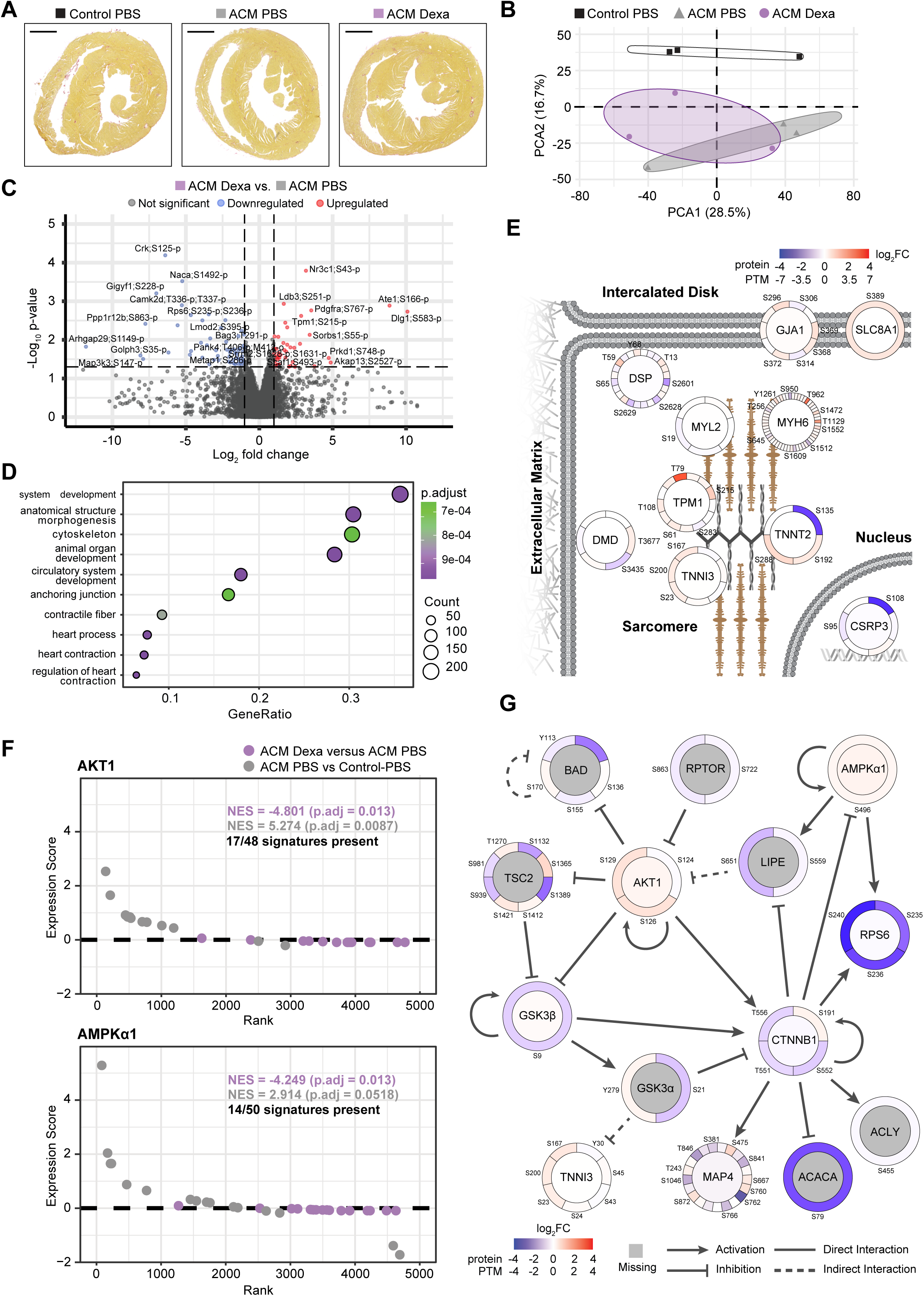
Dexamethasone treatment modulates cardiac contractility and Akt1/AMPKα1-signaling in an ACM mouse model. Proteomic and phospho-proteomic data of ventricular tissue from male ACM and Control mice 2 weeks post-induction, treated with PBS or Dexamethasone (Dexa) 1 week prior induction. Each dot represents the value of one animal. A) Representative images of picrosirius red collagen staining in heart sections at mid papillary level, scale bar = 1 mm. B) Principal component analysis (PCA) of phospho-proteomics data. Group identity ellipses drawn using Khachiyan algorithm. C) Volcano plot for all post-translational modifications (PTMs) detected in ACM PBS and ACM Dexa hearts. Labels indicate protein followed by the modified amino acid and its position and identity of the modification (p = phosphorylation, o = oxidation). One protein can have multiple PTMs. Log_2_ fold change (log_2_FC) threshold of ≤ −1 and ≥ 1, unadjusted p-value threshold of 0.05. Red dots indicate proteins upregulated in ACM Dexa mice compared to ACM PBS mice, blue dots indicate PTMs downregulated in ACM Dexa mice. D) Gene ontology (GO) analysis of proteins with at least 1 phospho-site with log_2_FC of ≤ −1 or ≥ 1 in ACM Dexa mice compared to Control PBS mice. Top 10 most significant terms (by adjusted p-value) are shown, including “biological process” and “cellular component” ontologies. E) Overview figure of approximate cellular locations of the 10 proteins that contribute to at least 9 of the top 10 most significant GO terms. The color of the inner circle represents log_2_FC of the protein, the color of the outer circle segments represents the log_2_FC of individual phosphosites. Red hues indicate upregulation in ACM Dexa mice, blue hues indicate downregulation. Phosphosites that occurred on multiple peptides in a protein were deduplicated as for PTM-SEA, but every instance of a PTM was kept rather than favoring the monophosphorylated version or the peptide with the fewest total PTMs. The mean of the log_2_FC for all PTMs was then used to obtain 1 log_2_FC value per phosphosite per protein and used for the plotting. F) Rank plot of kinases AKT1 and AMPKα1 for ACM Dexa versus ACM PBS and ACM PBS versus Control PBS comparisons following PTM-SEA analysis. Each PTM was deduplicated and an expression score calculated based on the q-value and log_2_FC (y-axis). Values were ranked low-to-high, with PTMs that occur in a specific kinase signature plotted (x-axis). Normalized enrichment score (NES) and accompanying adjusted p-value for each comparison are indicated. For AKT1, 17 PTMs out of 48 PTMs in the signature set associated with upregulation were present. For AMPKα1, 14 PTMs out of 50 PTMs in the signature set associated with upregulation were present. G) Interaction network drawn using Ingenuity Pathway Analysis (IPA) for AKT1, AMPKα1 and associated signature set proteins present in the PTM dataset. Circles representing protein and PTM log_2_FC are plotted as in E. Arrows indicate activation, blocked arrows inhibition. Full lines are direct interactions, dashed lines indirect interactions.

Initial principal component analysis of the total proteomics data revealed substantial overlap between the three conditions, with no clear separation either by genotype or treatment with few differentially expressed proteins between the conditions (Supp. Fig. 3A-C, Supp Table 4). Analysis of the phospho-proteome showed a distinction between control and mutant and control and dexamethasone treated animals, with multiple differentially phosphorylated proteins (Fig. 6B, Supp Fig. 3D, Supp Table 4).

To assess commonalities between the proteins bearing altered PTM levels in mutant mice treated with dexamethasone versus PBS, we performed GO term analysis on the proteins that had at least one differentially abundant PTM. We found that the top terms were related to cytoskeleton, contractile fibers and heart contraction, as well as anchoring junction and development (Fig. 6D, Supp. Table 5). Detailed analysis of the proteins associated with the respective terms revealed 10 proteins, which contribute to at least 9 of the top 10 most significant GO terms. Interestingly, these proteins are mainly located at the intercalated disc and the sarcomere contractile apparatus (Fig. 6E).

This suggests a contribution of cardiac contractility and junctional components to the effect of dexamethasone. However, this type of analysis collapses the available information down to protein level. It does not take into account that one protein might have multiple changes of PTM abundances in opposite directions, or that the downstream biological effect depends on specific sites or a combination of PTMs. To leverage this, we applied site-centric PTM Signature Enrichment Analysis (PTM-SEA^24^). After obtaining unique monophosphorylated peptides by de-duplication, this algorithm uses a curated database of site-specific kinase substrate signatures to identify alterations in kinase activity. This analysis revealed that the kinase substrate signatures depending on the activity of the kinases AKT1 and AMPKα1 were downregulated in ACM hearts treated with dexamethasone versus PBS treatment (Fig. 6F, Supp. Table 6). Moreover, these signatures were upregulated in ACM versus control animals indicating a revision of disease-caused alterations by dexamethasone treatment. Ingenuity Pathway Analysis (IPA^25^) of AKT1, AMPKα1 and associated kinase signature sets revealed a common network with major components of the pathways connected in a unified pathway, with AKT1, GSK3β and the junctional protein β-catenin (CTNN1) as central components (Fig. 6G).

In conclusion, treatment with dexamethasone reverted the PTM signature in ACM mouse hearts to the direction of healthy controls. Our data suggest a model in which AMPKα1/AKT1 kinase activity is modulated and interact with components of the canonical Wnt pathway, junctional proteins and the contractile apparatus leading to the protective effect of the pro-adhesive glucocorticoid dexamethasone.

## Discussion

In this study, we developed and validated a new screening method to analyze cell-cell adhesion in a large number of conditions. We applied this platform to evaluate an FDA-approved drug library under ACM-like conditions with depletion of the major desmosomal adhesion molecule DSG2. We identified a set of compounds strengthening cell-cell adhesion and validated the effect under conditions of disrupted adhesion by ACM-patient mutations. Analysis of the top pro-adhesive drugs revealed an enrichment of members of the glucocorticoid drug family and we selected dexamethasone as a representative for downstream experiments. Pilot *in vivo* experiments in an ACM mouse model showed a protective effect of the pro-adhesive dexamethasone with respect to cardiac output function and ECG changes. Mechanistically, we found changes in the phosphorylation state of the contractile and junctional apparatus and signaling via AKT1/AMPK to be potentially involved. In summary, this study established a novel high-throughput platform to evaluate cell-cell adhesion and applied it in the context of ACM. We identified glucocorticoids as pro-adhesive compounds with therapeutic potential *in vivo.* This can have important implications for the clinical context and future therapeutic strategies in ACM.

### Applications and limitations of a cell-cell adhesion-based screening

To our knowledge, we here report the first screening platform to evaluate cell-cell adhesion of eukaryotic cells on a large scale. This screen was conducted in a 2D setting with a well characterized cell line to reduce assay complexity. However, *in vivo* adhesion mostly depends on a 3D context, which is lacking in this system. We envision future development of the assay into 3D by involvement of tissue equivalents like organoids or assembloids e.g. derived from induced pluripotent stem cells.^30^ The presented screening results are further limited due to the employment of an epithelial cell line to model a cardiac disease. On the other hand, using this methodology the experiments have higher reproducibility, which is a prerequisite for screening approaches, yielding a large list of substances with the general potential to strengthen adhesion independent of the desmosomal function. However, validation of the results is needed to exclude cell type specific effects prior to downstream application of the compounds. With the general pipeline being established in this study, application to more complex conditions with other cell lines, cell types, and compound libraries is streamlined and can be performed in the future. Moreover, the application of the assay and implications of the revealed data are not limited to ACM but can be extended to the identification of adhesion modulators in the therapeutic context of other diseases with a patho-mechanistical role of defective cell-cell adhesion such as blistering skin diseases, inflammatory bowel disease, or tumor metastasis formation.^9–13^

### Defective cell-cell adhesion as therapeutic target in ACM

Defective desmosomal adhesion was shown to cause an ACM phenotype^4^ and our data confirmed a loss of cell-cell adhesion by expression of certain ACM patient mutations in different desmosomal components. This indicates loss of cell-cell adhesion as a common trunk in at least a subpopulation of ACM, independent from the gene affected by the respective mutation. Of note, stabilizing intercellular adhesion was shown to also enhance electrical coupling in an ACM model.^31^ This highlights strengthening of cell-cell adhesion as an interesting potentially therapeutic concept with its relevance not limited to patients with a single mutation but extended implications for a group of mutations and cases with gene-elusive ACM or variants of unknown significancy.

### Pro-adhesive effect of glucocorticoids

Based on the data of the screen, we identified the ability of glucocorticoids to strengthen intercellular adhesion when desmosomal function is impaired. Clinically, this drug class is mainly applied for anti-inflammatory therapy e.g. in autoimmune disease. Canonically, glucocorticoids bind to the glucocorticoid receptor (NR3C1), which translocates to the nucleus upon activation to regulate gene expression with different modes of action being described (e.g. direct binding to DNA, indirect modulation via NF-kappaB). In addition to this genomic effect, non-genomic activation is known involving signaling via second messengers and kinase pathways (eNOS, AKT1). Both mechanisms were described to repress inflammatory pathways and factors (e.g. COX-2, MAPKAP), which also includes adhesion molecules involved in immune cell adhesion and epithelial barrier function such as ICAM-1, E- and N-cadherins, DSG3 and different catenins.^32,33^

Based on the pro-adhesive effect, we performed here a first pilot *in vivo* experiment investigating the overall effect of glucocorticoids in the context of ACM by systematically applying a relatively high dose of dexamethasone with start of application prior to induction of the disease. Our data revealed a protective effect for several ACM characteristics under these conditions, which paves the way for detailed future analyses including onset of the therapy in later stages of the disease. While high-dose glucocorticoid shots are therapeutically used in the acute setting, long-term treatment is limited due to many side effects including muscle atrophy, insulin resistance and hypertension.^34,35^ Thus, evaluation of the relevance of glucocorticoids for ACM therapy is needed with respect to application of lower concentrations or compounds with improved pharmacological properties (e.g. PEGylation, targeted delivery via liposomes or nanoparticles) to reduce side effects. Importantly, we found a large range of concentrations to lead to a pro-adhesive effect *in vitro* making a protective effect possible also for lower concentrations.^34,36^ In addition, to lay the mechanistic basis for more targeted strategies reducing side effects, we analyzed the impact of dexamethasone treatment on the cardiac proteome and PTMs in mice prior to major structural changes. While the total protein levels remained largely unaltered in treated ACM hearts vs. control treatment, alterations in the phosphorylation state highlight components of the contractile apparatus and suppression of AKT1/AMPK signaling including downstream suppression of GSK3β and involvement of β-catenin. Interestingly, our data show that the dexamethasone effect was abolished in cells expressing a truncated version of JUP, a β-catenin analog. This suggests a functional contribution of JUP to the pro-adhesive effect. Moreover, the identified targets and pathways of dexamethasone are overlapping with targets and downstream factors of other compounds of the identified top 100 pro-adhesive drugs (e.g. PPARs, COX-2, AMPK). Interestingly, inhibition of GSK3β or modulation of PPARs was also shown to be protective in ACM models.^37–39^ Based on these data, a similar mechanism of the different compounds to strengthen cell-cell adhesion independent from the initial drug target can be envisioned with confirmatory studies needed in the future.

Our experiments identified a pro-adhesive effect of dexamethasone in cells expressing different ACM patient mutations, however, *in vivo* we cannot distinguish adhesion-dependent protective effects from anti-inflammatory effects or general cardioprotective effects of glucocorticoids.^35^ These effects can be beneficial in ACM additionally or independently from strengthening of adhesion, especially as inflammation and a myocarditis-like phenotype is described in subpopulations of ACM patients.^40,41^ Independent from the underlying mechanism, our data suggest a therapeutic potential for glucocorticoids in ACM. In line with this, case studies report a beneficial effect of glucocorticoid treatment in a child with inflammatory ACM and in ACM patients with DSP mutations.^42,43^ Thus, future evaluations are needed to clarify the mechanisms of the protective effect and clinical application of glucocorticoids for ACM therapy.

## Abbreviations

ACM: Arrhythmogenic Cardiomyopathy
DSG2: Desmoglein 2
DSC2: Desmocollin 2
JUP: Junctional Plakoglobin
DSP: Desmoplakin
PKP2: Plakophilin 2
WT: wildtype
KO: knock out
PTM: post-translational modification
F/R: forskolin/rolipram
FAC: Fractional area change
PVCs: premature ventricular contractions
ECG: Electrocardiogram
PTM-SEA: PTM Signature Enrichment Analysis
IPA: Ingenuity Pathway Analysis

## Acknowledgments

We thank Julian Zeiler (LMU Munich) for generation of the AsPC-1 DSG2 KO cells; Katja Gehmlich (University of Birmingham) for providing ACM mutant plasmids; Nicolas Schlegel (University of Würzburg) for providing the CaCo2 cells; Arnd Heuser (Animal Phenotyping Platform, Max-Delbrück Center, Berlin) for providing the *Dsg2-flox* mouse line; Judith Fülle and Christoph Ballestrem (University of Manchester) for providing the PKP2a construct; Aude Zimmermann and Lea Galvagno (DBM, University Basel) for technical assistance; Katarzyna Buczak (Proteomics Facility Basel, Biozentrum, University of Basel) for conduction of proteomic analysis; Alain Brühlhart and the team from the Animal Facility for animal care taking; Diego Calabrese (Histology Core Facility) and Evelina Bartoszek and Mike Abanto (Microscopy Core Facility), all DBM, University of Basel, Switzerland for support. Volker Spindler (University of Basel and UKE Hamburg) for proof reading of the manuscript and providing equipment.

## Author contributions

Conceptualization and funding: C. Schinner; Data acquisition: P. Hanns, R. Colpaert, R. Castellanos-Martinez, F. Weidner, S. Beensen, C. Schinner; Data Analysis: P. Hanns, R. Colpaert, R. Castellanos-Martinez, F. Weidner, F. Matthias, L. Xu, C. Schinner; Funding Acquisition: C. Schinner; Project Administration: P. Hanns, R. Colpaert, C. Schinner; Resources: C. Schinner; G. M. Kuster; Supervision: C. Schinner; Writing Original Draft Preparation: P. Hanns, R. Colpaert, C. Schinner; Writing - Review and Editing: all authors.

## Supplemental Material

Supplementary Tables 1–6

Supplementary Figure 1–3

## References

1. Corrado D, Link MS, Calkins H. Arrhythmogenic Right Ventricular Cardiomyopathy. N Engl J Med. 2017;376:61–72. doi: 10.1056/NEJMra1509267

2. Hoorntje ET, Te Rijdt WP, James CA, Pilichou K, Basso C, Judge DP, Bezzina CR, van Tintelen JP. Arrhythmogenic cardiomyopathy: pathology, genetics, and concepts in pathogenesis. Cardiovasc Res. 2017;113:1521–1531. doi: 10.1093/cvr/cvx150

3. Austin KM, Trembley MA, Chandler SF, Sanders SP, Saffitz JE, Abrams DJ, Pu WT. Molecular mechanisms of arrhythmogenic cardiomyopathy. Nat Rev Cardiol. 2019;16:519–537. doi: 10.1038/s41569-019-0200-7

4. Schinner C, Xu L, Franz H, Zimmermann A, Wanuske MT, Rathod M, Hanns P, Geier F, Pelczar P, Liang Y, et al. Defective Desmosomal Adhesion Causes Arrhythmogenic Cardiomyopathy by Involving an Integrin-alphaVbeta6/TGF-beta Signaling Cascade. Circulation. 2022;146:1610–1626. doi: 10.1161/CIRCULATIONAHA.121.057329

5. Schinner C, Vielmuth F, Rotzer V, Hiermaier M, Radeva MY, Co TK, Hartlieb E, Schmidt A, Imhof A, Messoudi A, et al. Adrenergic Signaling Strengthens Cardiac Myocyte Cohesion. Circ Res. 2017;120:1305–1317. doi: 10.1161/CIRCRESAHA.116.309631

6. Shoykhet M, Trenz S, Kempf E, Williams T, Gerull B, Schinner C, Yeruva S, Waschke J. Cardiomyocyte adhesion and hyperadhesion differentially require ERK1/2 and plakoglobin. JCI Insight. 2020;5. doi: 10.1172/jci.insight.140066

7. Schinner C, Erber BM, Yeruva S, Waschke J. Regulation of cardiac myocyte cohesion and gap junctions via desmosomal adhesion. Acta Physiol (Oxf). 2019;226:e13242. doi: 10.1111/apha.13242

8. Schinner C, Olivares-Florez S, Schlipp A, Trenz S, Feinendegen M, Flaswinkel H, Kempf E, Egu DT, Yeruva S, Waschke J. The inotropic agent digitoxin strengthens desmosomal adhesion in cardiac myocytes in an ERK1/2-dependent manner. Basic Res Cardiol. 2020;115:46. doi: 10.1007/s00395-020-0805-3

9. Dietrich N, Castellanos-Martinez R, Kemmling J, Heuser A, Schnoor M, Schinner C, Spindler V. Adhesion of pancreatic tumor cell clusters by desmosomal molecules enhances early liver metastases formation. Sci Rep. 2024;14:18189. doi: 10.1038/s41598-024-68493-6

10. Padmanaban V, Krol I, Suhail Y, Szczerba BM, Aceto N, Bader JS, Ewald AJ. E-cadherin is required for metastasis in multiple models of breast cancer. Nature. 2019;573:439–444. doi: 10.1038/s41586-019-1526-3

11. Kasperkiewicz M, Ellebrecht CT, Takahashi H, Yamagami J, Zillikens D, Payne AS, Amagai M. Pemphigus. Nat Rev Dis Primers. 2017;3:17026. doi: 10.1038/nrdp.2017.26

12. Schlegel N, Boerner K, Waschke J. Targeting desmosomal adhesion and signalling for intestinal barrier stabilization in inflammatory bowel diseases-Lessons from experimental models and patients. Acta Physiol (Oxf). 2021;231:e13492. doi: 10.1111/apha.13492

13. Aceto N, Bardia A, Miyamoto DT, Donaldson MC, Wittner BS, Spencer JA, Yu M, Pely A, Engstrom A, Zhu H, et al. Circulating tumor cell clusters are oligoclonal precursors of breast cancer metastasis. Cell. 2014;158:1110–1122. doi: 10.1016/j.cell.2014.07.013

14. Schlipp A, Schinner C, Spindler V, Vielmuth F, Gehmlich K, Syrris P, McKenna WJ, Dendorfer A, Hartlieb E, Waschke J. Desmoglein-2 interaction is crucial for cardiomyocyte cohesion and function. Cardiovasc Res. 2014;104:245–257. doi: 10.1093/cvr/cvu206

15. Gehmlich K, Syrris P, Peskett E, Evans A, Ehler E, Asimaki A, Anastasakis A, Tsatsopoulou A, Vouliotis AI, Stefanadis C, et al. Mechanistic insights into arrhythmogenic right ventricular cardiomyopathy caused by desmocollin-2 mutations. Cardiovasc Res. 2011;90:77–87. doi: 10.1093/cvr/cvq353

16. Fulle JB, Huppert H, Liebl D, Liu J, Alves de Almeida R, Yanes B, Wright GD, Lane EB, Garrod DR, Ballestrem C. Desmosome dualism - most of the junction is stable, but a plakophilin moiety is persistently dynamic. J Cell Sci. 2021;134. doi: 10.1242/jcs.258906

17. Wanuske MT, Brantschen D, Schinner C, Studle C, Walter E, Hiermaier M, Vielmuth F, Waschke J, Spindler V. Clustering of desmosomal cadherins by desmoplakin is essential for cell-cell adhesion. Acta Physiol (Oxf). 2021;231:e13609. doi: 10.1111/apha.13609

18. Bankhead P, Loughrey MB, Fernandez JA, Dombrowski Y, McArt DG, Dunne PD, McQuaid S, Gray RT, Murray LJ, Coleman HG, et al. QuPath: Open source software for digital pathology image analysis. Sci Rep. 2017;7:16878. doi: 10.1038/s41598-017-17204-5

19. Szklarczyk D, Santos A, von Mering C, Jensen LJ, Bork P, Kuhn M. STITCH 5: augmenting protein-chemical interaction networks with tissue and affinity data. Nucleic Acids Res. 2016;44:D380–384. doi: 10.1093/nar/gkv1277

20. Rimpler U. Funktionelle Charakterisierung von Desmocollin 2 während der Embryonalentwicklung und im adulten Herzen in der Maus. 2014.

21. Post H, Penning R, Fitzpatrick MA, Garrigues LB, Wu W, MacGillavry HD, Hoogenraad CC, Heck AJ, Altelaar AF. Robust, Sensitive, and Automated Phosphopeptide Enrichment Optimized for Low Sample Amounts Applied to Primary Hippocampal Neurons. J Proteome Res. 2017;16:728–737. doi: 10.1021/acs.jproteome.6b00753

22. Ahrne E, Glatter T, Vigano C, Schubert C, Nigg EA, Schmidt A. Evaluation and Improvement of Quantification Accuracy in Isobaric Mass Tag-Based Protein Quantification Experiments. J Proteome Res. 2016;15:2537–2547. doi: 10.1021/acs.jproteome.6b00066

23. Choi M, Chang CY, Clough T, Broudy D, Killeen T, MacLean B, Vitek O. MSstats: an R package for statistical analysis of quantitative mass spectrometry-based proteomic experiments. Bioinformatics. 2014;30:2524–2526. doi: 10.1093/bioinformatics/btu305

24. Krug K, Mertins P, Zhang B, Hornbeck P, Raju R, Ahmad R, Szucs M, Mundt F, Forestier D, Jane-Valbuena J, et al. A Curated Resource for Phosphosite-specific Signature Analysis. Mol Cell Proteomics. 2019;18:576–593. doi: 10.1074/mcp.TIR118.000943

25. Kramer A, Green J, Pollard J, Jr., Tugendreich S. Causal analysis approaches in Ingenuity Pathway Analysis. Bioinformatics. 2014;30:523–530. doi: 10.1093/bioinformatics/btt703

26. Kant S, Holthofer B, Magin TM, Krusche CA, Leube RE. Desmoglein 2-Dependent Arrhythmogenic Cardiomyopathy Is Caused by a Loss of Adhesive Function. Circ Cardiovasc Genet. 2015;8:553–563. doi: 10.1161/CIRCGENETICS.114.000974

27. Vielmuth F, Radeva MY, Yeruva S, Sigmund AM, Waschke J. cAMP: A master regulator of cadherin-mediated binding in endothelium, epithelium and myocardium. Acta Physiol (Oxf). 2023;238:e14006. doi: 10.1111/apha.14006

28. Spindler V, Waschke J. Desmosomal cadherins and signaling: lessons from autoimmune disease. Cell Commun Adhes. 2014;21:77–84. doi: 10.3109/15419061.2013.877000

29. Braga VM. Cell-cell adhesion and signalling. Curr Opin Cell Biol. 2002;14:546–556. doi: 10.1016/s0955-0674(02)00373-3

30. Zuppinger C. 3D Cardiac Cell Culture: A Critical Review of Current Technologies and Applications. Front Cardiovasc Med. 2019;6:87. doi: 10.3389/fcvm.2019.00087

31. Schinner C, Erber BM, Yeruva S, Schlipp A, Rotzer V, Kempf E, Kant S, Leube RE, Mueller TD, Waschke J. Stabilization of desmoglein-2 binding rescues arrhythmia in arrhythmogenic cardiomyopathy. JCI Insight. 2020;5. doi: 10.1172/jci.insight.130141

32. Carayol N, Campbell A, Vachier I, Mainprice B, Bousquet J, Godard P, Chanez P. Modulation of cadherin and catenins expression by tumor necrosis factor-alpha and dexamethasone in human bronchial epithelial cells. Am J Respir Cell Mol Biol. 2002;26:341–347. doi: 10.1165/ajrcmb.26.3.4684

33. Mao X, Cho MJT, Ellebrecht CT, Mukherjee EM, Payne AS. Stat3 regulates desmoglein 3 transcription in epithelial keratinocytes. JCI Insight. 2017;2. doi: 10.1172/jci.insight.92253

34. Vandewalle J, Luypaert A, De Bosscher K, Libert C. Therapeutic Mechanisms of Glucocorticoids. Trends Endocrinol Metab. 2018;29:42–54. doi: 10.1016/j.tem.2017.10.010

35. Hafezi-Moghadam A, Simoncini T, Yang Z, Limbourg FP, Plumier JC, Rebsamen MC, Hsieh CM, Chui DS, Thomas KL, Prorock AJ, et al. Acute cardiovascular protective effects of corticosteroids are mediated by non-transcriptional activation of endothelial nitric oxide synthase. Nat Med. 2002;8:473–479. doi: 10.1038/nm0502-473

36. Rhen T, Cidlowski JA. Antiinflammatory action of glucocorticoids--new mechanisms for old drugs. N Engl J Med. 2005;353:1711–1723. doi: 10.1056/NEJMra050541

37. Chelko SP, Asimaki A, Andersen P, Bedja D, Amat-Alarcon N, DeMazumder D, Jasti R, MacRae CA, Leber R, Kleber AG, et al. Central role for GSK3beta in the pathogenesis of arrhythmogenic cardiomyopathy. JCI Insight. 2016;1. doi: 10.1172/jci.insight.85923

38. Qiu Z, Zhao Y, Tao T, Guo W, Liu R, Huang J, Xu G. Activation of PPARalpha Ameliorates Cardiac Fibrosis in Dsg2-Deficient Arrhythmogenic Cardiomyopathy. Cells. 2022;11. doi: 10.3390/cells11203184

39. Reisqs JB, Moreau A, Charrabi A, Sleiman Y, Meli AC, Millat G, Briand V, Beauverger P, Richard S, Chevalier P. The PPARgamma pathway determines electrophysiological remodelling and arrhythmia risks in DSC2 arrhythmogenic cardiomyopathy. Clin Transl Med. 2022;12:e748. doi: 10.1002/ctm2.748

40. Lin YN, Ibrahim A, Marban E, Cingolani E. Pathogenesis of arrhythmogenic cardiomyopathy: role of inflammation. Basic Res Cardiol. 2021;116:39. doi: 10.1007/s00395-021-00877-5

41. Chelko SP, Penna VR, Engel M, Shiel EA, Centner AM, Farra W, Cannon EN, Landim-Vieira M, Schaible N, Lavine K, et al. NFkB signaling drives myocardial injury via CCR2+ macrophages in a preclinical model of arrhythmogenic cardiomyopathy. J Clin Invest. 2024;134. doi: 10.1172/JCI172014

42. Peretto G, Barzaghi F, Cicalese MP, Di Resta C, Slavich M, Benedetti S, Giangiobbe S, Rizzo S, Palmisano A, Esposito A, et al. Immunosuppressive therapy in childhood-onset arrhythmogenic inflammatory cardiomyopathy. Pacing Clin Electrophysiol. 2021;44:552–556. doi: 10.1111/pace.14153

43. Gasperetti A, Carrick RT, Muller S, Murray B, Adamo L, Bauce B, McNally E, Helms A. Desmoplakin Cardiomyopathy: Role of Inflammation and Potential Role of Disease-Modifying Therapies. Curr Cardiol Rep. 2025;27:12. doi: 10.1007/s11886-024-02183-7

44. Delva E, Tucker DK, Kowalczyk AP. The desmosome. Cold Spring Harb Perspect Biol. 2009;1:a002543. doi: 10.1101/cshperspect.a002543

