## Supplemental Material for "Screening for pro-adhesive compounds and their relevance as therapeutic approach in Arrhythmogenic Cardiomyopathy"

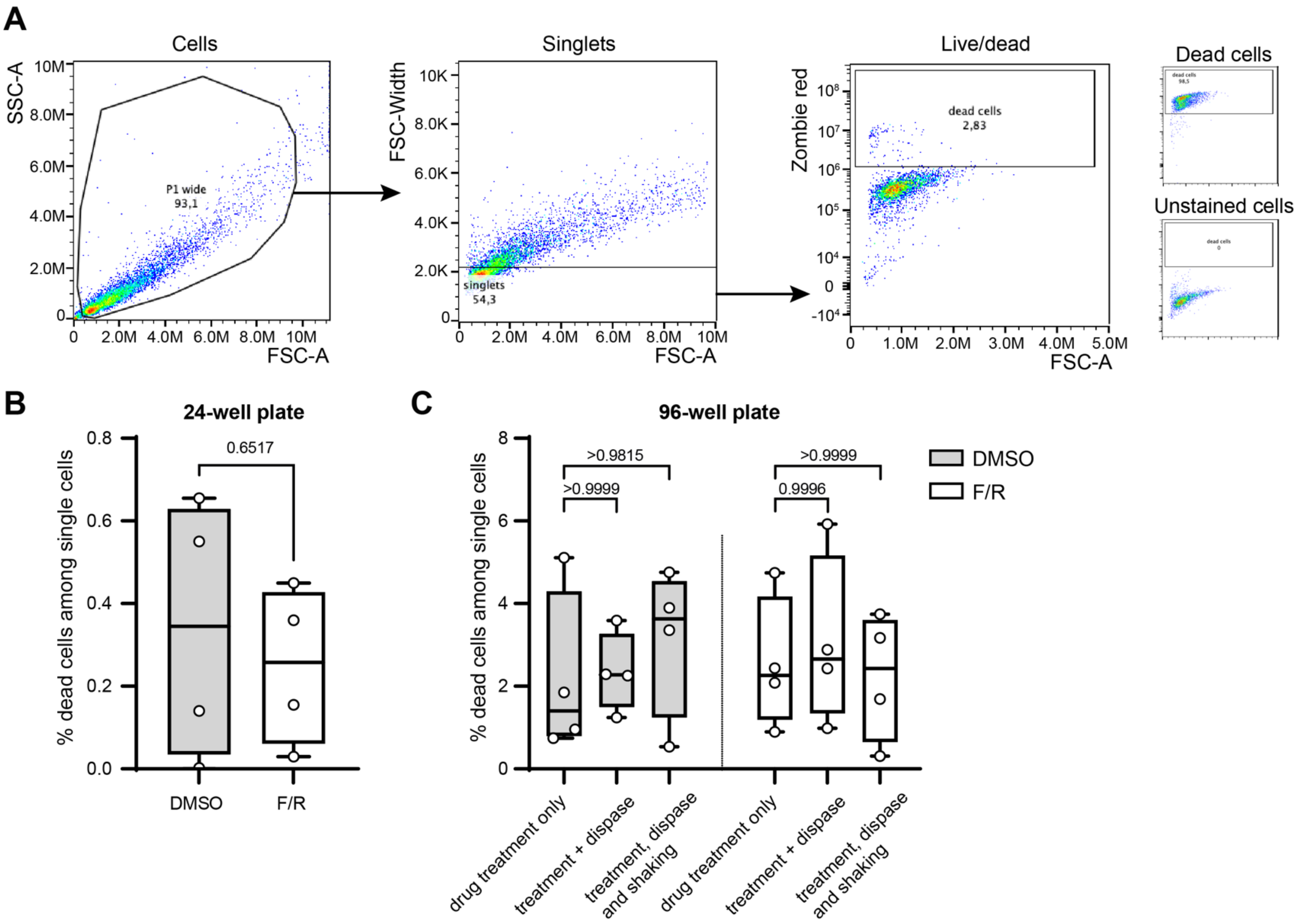

**Supplementary Figure 1. Cell viability is not affected during cell-cell adhesion assays.** A) Flow cytometry gating strategy to assess cell viability. Cells were gated for particle size by SSC-A vs. FSC-A, followed by single cell gating by FSC-Width vs. FSC-A. Zombie Red staining was performed to determine the dead cell population. Unstained cells and dead cells (heated to 95°C) served as negative and positive control, respectively, to set the "dead cells" gate. B) Cell viability analysis in 24-well plate after DMSO or forskolin/rolipram (F/R) treatment. Each dot represents a biological replicate (independent cell seeding) as mean of 4 technical replicates, unpaired t-test. C) Cell viability analysis in 96-well plate after DMSO or F/R treatment with collection of samples prior and after dispase-based cell detachment, and after application of shear stress to the respective cell monolayer. Each dot represents a biological replicate (independent cell seeding) as mean of 4 technical replicates, ordinary one-way ANOVA with Sidak post-hoc test. $p>0.95$ for DMSO versus F/R for the respective treatment (not indicated).

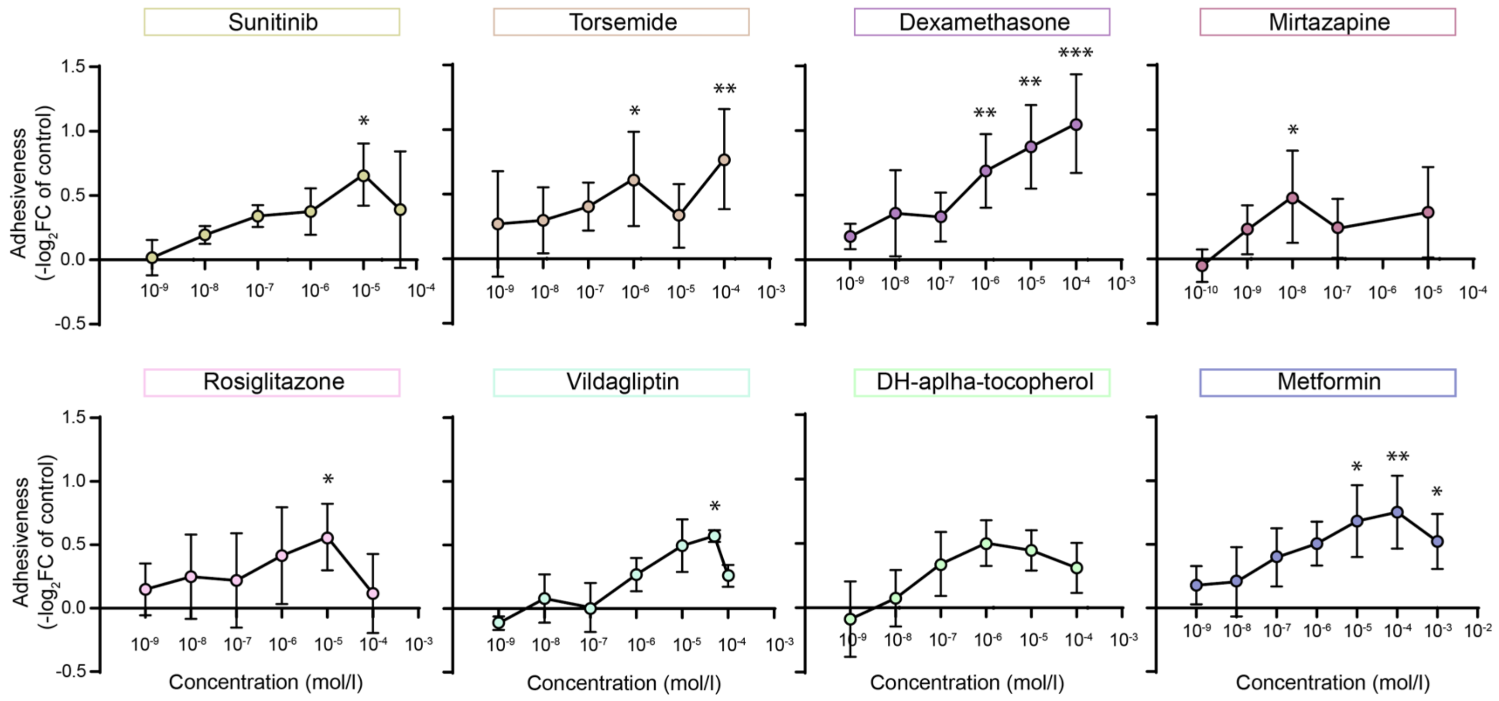

**Supplementary Figure 2. Dose-response curve of selected pro-adhesive compounds.** Dose-response curves of selected pro-adhesive compounds tested in CaCo2 cells overexpressing DSG2-W2A. Dots are indicating the mean of 3 - 6 biological replicates (independent cell seedings), with an average of 4 technical replicates, error bars represent the standard deviation. Ordinary one-way ANOVA with Dunnett post-hoc multiple comparison test or Kruskal-Wallis test with Dunn's correction vs. corresponding control condition was applied. $^{*}p < 0.05$, $^{**}p < 0.01$, $^{***}p < 0.001$.

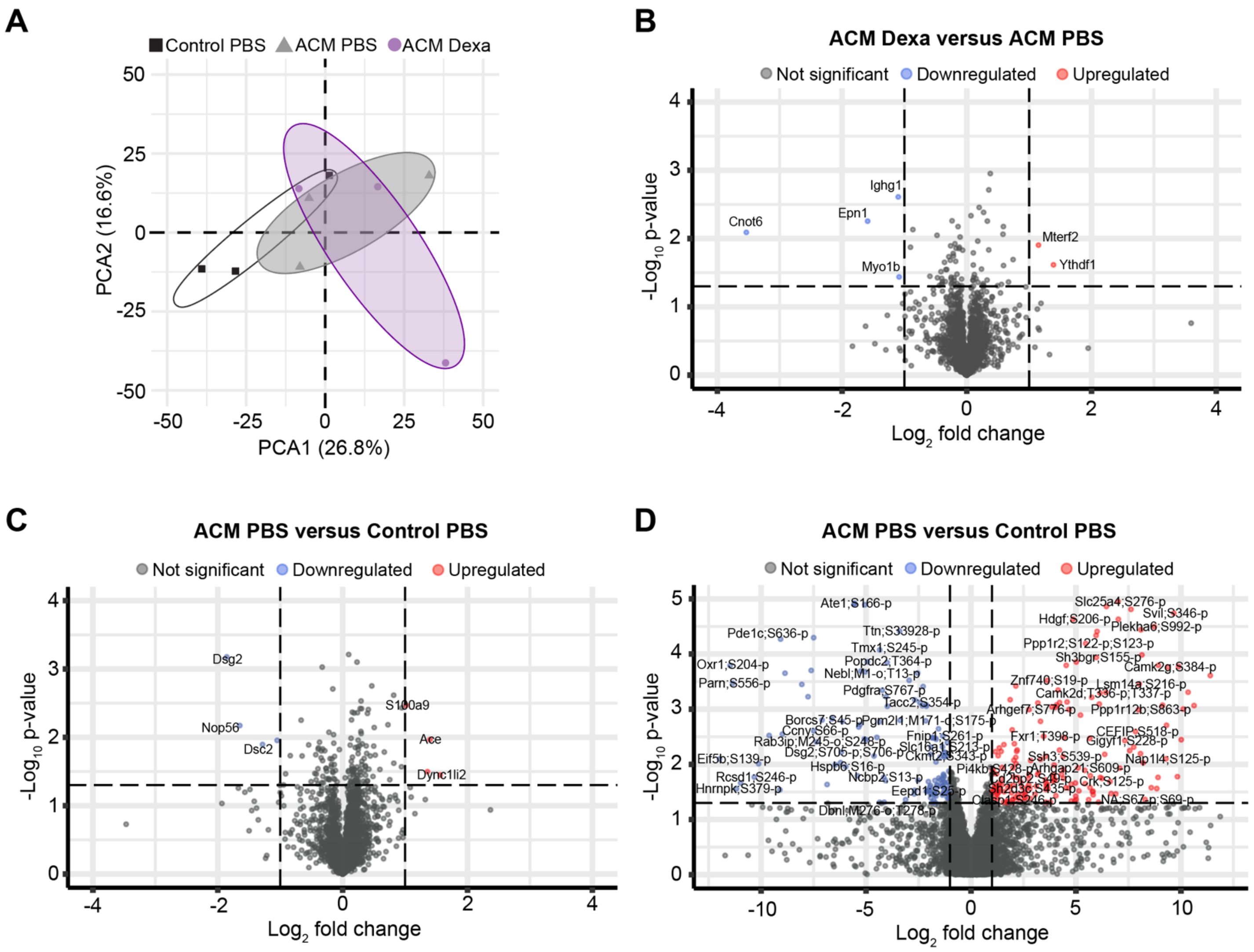

**Supplementary Figure 3. Extended proteomic and phospho-proteomic analyses.** A) Principal component analysis (PCA) of proteomics data. Group identity ellipses drawn using Khachiyan algorithm. B, C) Volcano plot of all proteins detected in dexamethasone treated ACM (ACM Dexa) versus PBS treated ACM animals (ACM PBS) and ACM PBS versus PBS treated control mice (Control PBS). $Log_2$fold change threshold of ±1, unadjusted p-value threshold of 0.05. D) Volcano plot for all post-translational modifications (PTMs) detected in ACM PBS and Control PBS mice. Labels indicate protein followed by the modified amino acid and its position and identity of the modification (p = phosphorylation, o = oxidation). One protein can have multiple PTMs. $Log_2$ fold change threshold of ≤ -1 and ≥ 1, unadjusted p-value threshold of 0.05.

| Gene | Variation | Classification according to ClinVar | Identifiers | Variation ID | Accession number |
|---|---|---|---|---|---|
| DSG2 | W51S | likely pathogenic | NM_001943.5(DSG2):c.152G>C (p.Trp51Ser) | 199796 | VCV000199796.2 |
| | W51R | unknown significance | NM_001943.5(DSG2):c.151T>C (p.Trp51Arg) | 420241 | VCV000420241.1 |
| | N266S | pathogenic | NM_001943.5(DSG2):c.797A>G (p.Asn266Ser) | 16815 | VCV000016815.1 |
| | A517V | conflicting (unknown significance/likely benign/benign) | NM_001943.5(DSG2):c.1550C>T (p.Ala517Val) | 163210 | VCV000163210.33 |
| | Q558* | pathogenic | NM_001943.5(DSG2):c.1672C>T (p.Gln558Ter) | 1075931 | VCV001075931.8 |
| | G812S | conflicting (likely pathogenic/unknown significance) | NM_001943.5(DSG2):c.2434G>A (p.Gly812Ser) | 199817 | VCV000199817.52 |
| | Q993H | likely pathogenic | NM_001943.5(DSG2):c.2979G>T (p.Gln993His) | 996557 | VCV000996557.1 |
| DSC2 | R203C | unknown significance | NM_024422.6(DSC2):c.607C>T (p.Arg203Cys) | 72377 | VCV000072377.18 |
| | F250S | likely pathogenic | NM_024422.6(DSC2):c.749T>C (p.Phe250Ser) | 523128 | VCV000523128.1 |
| | T275M | conflicting (likely pathogenic/unknown significance) | NM_024422.6(DSC2):c.824C>T (p.Thr275Met) | 222557 | VCV000222557.10 |
| | R375G | unknown significance | NM_024422.6(DSC2):c.1123C>G (p.Arg375Gly) | 199782 | VCV000199782.18 |
| | Q554* | pathogenic | NM_024422.6(DSC2):c.1660C>T (p.Gln554Ter) | 235909 | VCV000235909.14 |
| JUP | N638H | conflicting (unknown significance/likely benign) | NM_002230.4(JUP):c.1912A>C (p.Asn638His) | 323165 | VCV000323165.15 |
| | W680* | pathogenic | NM_002230.4(JUP):c.2039G>A (p.Trp680Ter) | 1363212 | VCV001363212.5 |
| DSP | T564I | pathogenic | NM_004415.4(DSP):c.1691C>T (p.Thr564Ile) | 157673 | VCV000157673.5 |
| PKP2 | V406fs | pathogenic | NM_001005242.3(PKP2):c.1211dup (p.Val406fs) | 45015 | VCV000045015.41 |
| | K472R | unknown significance | NM_001005242.3(PKP2):c.1415A>G (p.Lys472Arg) | 201985 | VCV000201985.17 |
| | S571F | pathogenic | NM_001005242.3(PKP2):c.1712C>T (p.Ser571Phe) | 406555 | VCV000406555.7 |
| | R691* | pathogenic | NM_001005242.3(PKP2):c.2071C>T (p.Arg691Ter) | 6755 | VCV000006755.40 |

**Supplementary Table 1. Additional information for patient derived variants applied in Figure 1.**

Classification according to ClinVar data base and respective variant IDs as indicated.

**Supplementary Table 2. Compiled results of the adhesion-based drug library screen.** Data are presented as adhesiveness defined as $-\log_2 FC$ of the revealed number of fragments normalized to the respective vehicle control. Drugs are ordered by decreasing maximal adhesiveness of the tested concentrations. Additional information of drug pathways and targets are based on the information provided by MedChemExpress (MCE).

| | | Control PBS | | Control Dexamethasone | | ACM PBS | | ACM Dexamethasone | |
|---|---|---|---|---|---|---|---|---|---|
| *Parameter* | *Unit* | *Mean* | *SD* | *Mean* | *SD* | *Mean* | *SD* | *Mean* | *SD* |
| **Body weight (8 weeks)** | g | 26.96 | 6.27 | 23.50 | 4.98 | 25.23 | 4.72 | 27.16 | 5.43 |
| **ECG (baseline)** | | | | | | | | | |
| **Heart Rate** | BPM | 435.71 | 27.56 | 492.80 | 51.95 | 508.60* | 52.38 | 481.96 | 57.46 |
| **PR Interval** | sec | 0.04 | 0.01 | 0.04 | 0.00 | 0.04 | 0.00 | 0.04 | 0.00 |
| **P Duration** | sec | 0.01 | 0.00 | 0.01 | 0.00 | 0.01 | 0.00 | 0.01 | 0.00 |
| **QRS Interval** | sec | 0.01 | 0.01 | 0.01 | 0.00 | 0.01 | 0.00 | 0.01 | 0.00 |
| **P Amplitude** | mV | 0.09 | 0.03 | 0.10 | 0.00 | 0.09 | 0.03 | 0.12 | 0.03 |
| **Q Amplitude** | mV | -0.01 | 0.01 | 0.00 | 0.00 | -0.02 | 0.01 | -0.03 | 0.02 |
| **R Amplitude** | mV | 1.27 | 0.21 | 1.17 | 0.29 | 1.35 | 0.40 | 1.47 | 0.40 |
| **S Amplitude** | mV | -0.36 | 0.12 | -0.43 | 0.14 | -0.37 | 0.17 | -0.44 | 0.27 |
| **J Amplitude** | mV | 0.06 | 0.08 | 0.17 | 0.13 | 0.06 | 0.06 | 0.06 | 0.08 |
| **QRS Amplitude** | mV | 1.63 | 0.19 | 1.60 | 0.42 | 1.72 | 0.51 | 1.91 | 0.54 |
| **ECG (4 weeks)** | | | | | | | | | |
| **Heart Rate** | BPM | 435.13 | 40.87 | 468.94 | 34.87 | 491.13 | 52.85 | 462.76 | 46.78 |
| **PR Interval** | sec | 0.04 | 0.01 | 0.04 | 0.00 | 0.04 | 0.00 | 0.04 | 0.00 |
| **P Duration** | sec | 0.01 | 0.00 | 0.01 | 0.00 | 0.01 | 0.00 | 0.01 | 0.00 |
| **QRS Interval** | sec | 0.01 | 0.00 | 0.01 | 0.00 | 0.01 | 0.00 | 0.01 | 0.00 |
| **P Amplitude** | mV | 0.08 | 0.03 | 0.10 | 0.02 | 0.09 | 0.02 | 0.11 | 0.03 |
| **Q Amplitude** | mV | -0.01 | 0.02 | 0.01 | 0.00 | -0.02 | 0.02 | -0.02 | 0.02 |
| **R Amplitude** | mV | 1.25 | 0.26 | 1.37 | 0.30 | 1.38 | 0.50 | 1.52 | 0.51 |
| **S Amplitude** | mV | -0.40 | 0.15 | -0.56 | 0.17 | -0.36 | 0.18 | -0.35 | 0.26 |
| **J Amplitude** | mV | 0.02 | 0.08 | 0.10 | 0.10 | 0.05 | 0.06 | 0.09 | 0.10 |
| **QRS Amplitude** | mV | 1.65 | 0.27 | 1.94 | 0.46 | 1.74 | 0.62 | 1.87 | 0.68 |
| **ECG (8 weeks)** | | | | | | | | | |
| **Heart Rate** | BPM | 477.01 | 65.11 | 483.12 | 43.24 | 509.53 | 57.24 | 486.65 | 59.50 |
| **PR Interval** | sec | 0.04 | 0.00 | 0.04 | 0.00 | 0.04 | 0.00 | 0.04 | 0.00 |
| **P Duration** | sec | 0.01 | 0.00 | 0.01 | 0.00 | 0.01 | 0.00 | 0.01 | 0.00 |
| **QRS Interval** | sec | 0.01 | 0.00 | 0.01 | 0.00 | 0.01 | 0.00 | 0.01 | 0.00 |
| **P Amplitude** | mV | 0.10 | 0.02 | 0.10 | 0.02 | 0.10 | 0.02 | 0.10 | 0.03 |
| **Q Amplitude** | mV | -0.01 | 0.02 | 0.00 | 0.01 | -0.01 | 0.01 | -0.02 | 0.03 |
| **R Amplitude** | mV | 1.19 | 0.23 | 1.22 | 0.33 | 1.03 | 0.32 | 1.40 | 0.43 |
| **S Amplitude** | mV | -0.37 | 0.09 | -0.40 | 0.16 | -0.14* | 0.13 | -0.20 | 0.24 |
| **J Amplitude** | mV | 0.03 | 0.10 | 0.15 | 0.08 | 0.08 | 0.11 | 0.06 | 0.08 |
| **QRS Amplitude** | mV | 1.56 | 0.28 | 1.62 | 0.46 | 1.17 | 0.37 | 1.60 | 0.55 |
| **Echocardiography, SAX, B-mode (8 weeks)** | | | | | | | | | |
| **RV Area;s** | $mm^2$ | 3.31 | 0.79 | 2.63 | 1.12 | 4.38 | 1.24 | $2.53^{\#}$ | 0.71 |
| **RV Area;d** | $mm^2$ | 5.51 | 1.51 | 4.46 | 1.41 | 5.57 | 1.40 | 4.35 | 0.98 |
| **FAC** | % | 38.72 | 11.42 | 41.87 | 12.93 | 21.53* | 8.03 | $42.07^{\#}$ | 9.70 |
| **Echocardiography, SAX, LV trace M-mode (8 weeks)** | | | | | | | | | |
| **Heart Rate** | BPM | 458.86 | 47.07 | 470.25 | 29.83 | 545.09* | 55.44 | 526.08 | 56.41 |
| **Diameter;s** | mm | 3.05 | 0.43 | 2.64 | 0.21 | 2.78 | 0.50 | 2.63 | 0.49 |
| **Diameter;d** | mm | 4.23 | 0.36 | 3.97 | 0.52 | 3.93 | 0.45 | 3.85 | 0.39 |
| **Volume;s** | µl | 37.53 | 11.90 | 25.92 | 5.40 | 30.38 | 13.59 | 26.78 | 11.81 |
| **Volume;d** | µl | 80.57 | 15.44 | 69.94 | 23.03 | 68.54 | 18.93 | 64.72 | 14.95 |
| **Stroke volume** | µl | 43.05 | 5.51 | 44.03 | 17.76 | 38.16 | 8.34 | 37.93 | 6.09 |
| **Ejection fraction** | % | 54.43 | 7.52 | 61.94 | 4.64 | 56.99 | 9.69 | 60.25 | 9.81 |
| **Fractional shortening** | % | 28.08 | 4.73 | 32.97 | 3.66 | 29.80 | 6.66 | 32.01 | 6.56 |
| **Cardiac Output** | ml/min | 19.73 | 3.11 | 20.90 | 9.39 | 20.73 | 4.74 | 19.70 | 2.84 |
| **LV mass** | mg | 113.68 | 19.31 | 113.11 | 43.37 | 119.81 | 19.49 | 105.91 | 21.60 |
| **LV mass Cor** | mg | 90.94 | 15.45 | 90.49 | 34.70 | 95.84 | 15.59 | 84.73 | 17.28 |
| **LVAW;s** | mm | 1.12 | 0.17 | 1.21 | 0.23 | 1.23 | 0.16 | 1.18 | 0.14 |
| **LVAW;d** | mm | 0.78 | 0.11 | 0.81 | 0.12 | 0.89 | 0.08 | 0.83 | 0.12 |
| **LAPW;s** | mm | 0.97 | 0.13 | 1.09 | 0.15 | 1.16 | 0.24 | 1.13 | 0.16 |
| **LAPW;d** | mm | 0.67 | 0.09 | 0.72 | 0.10 | 0.76 | 0.14 | 0.71 | 0.10 |

**Supplementary Table 3. ECG and echocardiography measurements with selected data presented in Figure 5.** Mean values with respective standard deviation (SD) of acquired parameters for same animals presented in figure 5. ECG data: Two-way RM ANOVA with Dunnett's post hoc test, repeated measurements of same animal matched. *: $p<0.05$ vs. Control PBS; #: $p<0.05$ vs. ACM PBS. Echocardiography data acquired in short axis view (SAX) at mid-papillary level, right ventricle (RV), left ventricle (LV), end-systolic (s) and diastolic (d) measurement, anterior wall (AW), posterior wall (PW). For RV Area;d, a Kruskal-Wallis test with Dunn's post hoc test was performed, all other panels: ordinary one-way ANOVA with Sidak's post hoc test.

**Supplementary Table 4. Proteomic and phosphoproteomic data and analysis results.** Sheet 1) $Log_2$ transformed intensities per detected protein for each sample, as well as the p-value, adjusted p-value and $Log_2$ fold change ($log_2FC$) for each comparison between conditions. T-test with Benjamini and Hochberg adjustment. Sheet 2) $Log_2$ transformed intensities per detected phosphorylated peptide for each sample, as well as the coefficient of variation (cv), fraction of NA features that were assigned a pseudocount (fracNAFeatures), p-value and adjusted p-value for each comparison between conditions. T-test with Benjamini and Hochberg adjustment.

**Supplementary Table 5. Gene Ontology analysis of phosphoproteomics data.** Gene ontology analysis on the proteins that had at least one differentially abundant PTM. Results for each comparison between conditions are shown on separate sheets. One-sided Fisher's exact test with Benjamini and Hochberg adjustment.

**Supplementary Table 6. PTM-SEA input data and analysis results.** Sheet 1) Calculated peptide_score for each detected phosphorylation site following deduplication for each comparison between conditions. Data in GCT 1.3 format. Sheet 2) Results of PTM-SEA analysis. For each comparison between conditions, per signature, the p-value, adjusted p-value and normalized enrichment score are shown. Statistical test as implemented in the original paper[24], with Benjamini and Hochberg adjustment. Data in GCT 1.3 format.
